# Hypoxia-induced extracellular matrix changes are conserved in cancer and directly impact radiotherapy benefit

**DOI:** 10.1101/2025.04.04.647191

**Authors:** C. Guerrero Quiles, A. Julia Gonzalez, A. S. Foussat, A. Olympitis, T. Lodhi, V. Smith, M. Reardon, Rekaya Shabbir, S. Lunj, K. Reeves, A. Baker, Michael Eyers, Gayle Marshall, T.A.D. Smith, Peter Hoskin, Nicholas D. James, Emma Hall, Robert A. Huddart, Nuria Porta, V. Biolatti, J. D. Humphries, M. J. Humphries, A. Choudhury, C.M. West

## Abstract

Our overarching aim was to determine how hypoxia affects the extracellular matrix (ECM). Transcriptomic analysis (21,941 patients; 10 cancer types) identified ECM remodelling as the predominant pathway affected by hypoxia. Multi-omics confirmed that hypoxia impacts ECM organisation and collagen degradation; 53 ECM genes were affected, of which 74% were HIF1/HIF2-regulated. Spatial transcriptomics highlighted different hypoxia remodelling processes in tumour and stroma. Five ECM genes commonly affected in tumour and *in vitro* constituted a signature. This signature was independently prognostic and independently predictive of radiotherapy benefit in multiple malignancies. Patients within either high or low hypoxic-ECM score tertiles benefited from radiotherapy versus surgery. Hypoxic ECMs generated *in vitro* increased adhesion and decreased migration of cancer cells, an effect enhanced by irradiation. Immunofluorescence demonstrated that hypoxia decreased collagen fibre number, and irradiation decreased cell-ECM interactions. Taken together, these findings demonstrate hypoxia induces pan-cancer ECM changes, directly impacting radiotherapy benefit.

## Introduction

Hypoxia (<2% O_2_) is a feature of solid tumours and associates with a poor prognosis^1–3^. Cellular responses to hypoxia are primarily mediated by hypoxia-inducible factors (HIFs)^4^, although severe hypoxia (<0.2% O_2_) triggers HIF-independent mechanisms such as unfolded protein (UPR) and DNA damage (DDR) responses^5^. These mechanisms alter the expression of 1-1.5% of the human genome^6^, leading to metabolic switching, epithelial-mesenchymal-transition (EMT), and genomic instability, all increasing tumour aggressiveness^7^. Beyond these pathways, hypoxia remodelling of the extracellular matrix (ECM) is emerging as a regulator of treatment resistance and metastatic spread^1^, but is less widely studied.

The ECM is a complex network of proteins, including proteoglycans (e.g., fibronectin [FN1]), collagens [COLs], enzymes (e.g., metalloproteinase [MMPs]), and soluble factors (e.g., transforming growth factor β [TGF-β])^8^. The ECM interacts with cytoskeletal and focal adhesion (FA) proteins via cell-surface integrins [ITGs], activating phosphorylation cascades (e.g., FA kinase [FAK]) through vinculin (VCL) and paxillin (PXN) signalling^9–11^. FAs regulate cell/ECM interactions, affecting cellular processes, such as EMT and cadherin signalling^11^. Crosstalk between cadherins and ITGs further regulates FA formation and signalling^11^. During cancer development, the ECM undergoes intense remodelling, acquiring a pro-cancerous fibrotic phenotype^12^. Gilkes *et al*. review proposed hypoxia further enhances ECM remodelling and fibrosis - processes driven by HIF activation of intracellular collagen-modifying enzymes that promoted metastasis and tumour development^10^.

There are important interactions between the ECM and radiotherapy. ECM stiffness due to fibrosis activates FAK signalling through FAs, inducing radioresistance^13,14^. Furthermore, radiotherapy also modulates cadherin and FA levels, enhancing EMT^15^. However, the hypoxic ECM in cancer remains poorly characterised, with contradictory reports in the literature^16–18^. Here, we comprehensively characterised the hypoxia-associated ECM at a pan-cancer level. We derived a hypoxic-ECM signature prognostic and predictive of radiotherapy benefit in multiple cancer types.

## Materials and methods

### Cohorts

Supplementary Table 1 summarises patient cohorts. Cohorts (n=131) with transcriptomic data were downloaded for 10 cancer types with outcome data when available. Three bladder cancer cohorts available at our institution were also used (BCON, BC2001, Christie GemX). BCON and BC2001 transcriptomic data were previously generated^19,20^. Christie GemX (09/H1013/24) is an unpublished muscle-invasive bladder cancer (MIBC) cohort (n=184) described in Supplementary methods.

### Meta-analysis of transcriptomic data

Published hypoxia-associated cancer gene signatures were available for: bladder^19^, breast^21^, cervix^22^, colorectal^23^, brain^24^, liver^25^, head & neck^26^, lung^27^, pancreatic^28^, prostate^29^, and sarcoma^30^. Tumours were classified into high and low hypoxia-scores as described in the publications. For cervix, brain and pancreas signatures, patients were median-dichotomised into high and low based on their hypoxia-scores. Transcriptomic data was normalised and log2 transformed using *DESeq2:vst* (v.1.44.0)^31^. *Limma* (v3.60.3)^32^ and *qvalue* (v.3.20)^33^ calculated fold changes (FCs), p-values, adjusted p-values, q-values and FDR independently for each cohort. Significant genes (FDR<0.05, FC>1 or FC<-1) were identified and data integrated using random effects modelling (*metafor:rma* package)^34^. Cohorts with <20 differentially expressed genes (DEGs) were excluded (n=37 cohorts, Supplementary Table 1). Details are described in Supplementary Methods. ECM ontology was performed using The Matrisome Project Database^35^ as reference.

### *In vitro* multi-omics and mechanistic studies

Procedures are described in Supplementary Methods. Bladder cancer cells (T24, UMUC3, J82, RT4) were grown for 24 h (transcriptomics, ChIPseq) or 7 days (proteomics) under normoxia (20.9% O_2_) or hypoxia (0.2% O_2_). Omic data were generated and analysed following previously described protocols^36–40^. Western Blotting measured SLC2A1 protein expression. Attachment and scratch assays measured cell adhesion and migration for irradiated cells (0-8 Gy) seeded onto normoxic (20.9% O_2_) or hypoxic (0.2% O_2_) cell-derived ECMs (CDMs). Immunofluorescence measured FN1, COL5 and COL1 total levels, fibre numbers, morphology, and the fibres co-localisation with N-cadherin, E-cadherin, and VCL/PXN.

### Digital spatial profiling (DSP)

Formalin-fixed, paraffin-embedded (FFPE) diagnostic blocks were available from BCON trial patients^41^ under ethics license (LREC 09/H1013/24). Tissue microarrays (TMAs) produced for other studies^42,43^ underwent full transcriptomic DSP as described previously^44^. TMAs were stained for CA9 and cytokeratin. Regions of interest (ROIs) were outlined in tumour (n=47 ROIs) and stroma (n=32 ROIs) in samples from 28 patients. Genes were normalised and those undetected in ≤10% of ROIs discarded. ECM genes were annotated using The Matrisome Project Database^35^. Gene expression scores were calculated as the hypoxia transcriptomic meta-analysis, excluding gene frequency variable. *In silico* analyses were performed using R (v.4.4.2) packages: *org*.*Hs*.*eg*.*db, AnnotationDbi, ClusterProfiler, DOSE, ggplot2, enrichplot; ReactomePA; GoSemSIM; ggVennDiagram*.

### Analysis of clinical cohorts

Supplementary Methods and Table 1 summarise clinical endpoints. Patients were tertile-stratified into low (≤32%) medium (33 – 65%) and high (≥66%) based on their hypoxic-ECM scores for each cohort. Each malignancy was analysed as a single cohort. We assessed predictive capacity (radiotherapy versus surgery) by meta-analysing datasets (bladder and head & neck cancer). Endpoint event estimates were calculated using Kaplan–Meier (KM) and Cox (univariable and multivariable) regression. Analyses were performed using R (v.4.3.0) packages: *tibble, survival, survminer*. For bladder cancer, a prediction model was constructed using artificial neural networks (ANN) as described in the Supplementary Methods.

## Results

### Hypoxia affects the expression of ECM-related genes at a pan-cancer level

Random-effects modelling identified DEGs in hypoxia-high and -low tumours (10 cancer types; 131 cohorts; 21,941 patients). We identified 954 pan-cancer DEGs, 17.3% being ECM genes. Classic hypoxia markers e.g. CA9 and SLC2A1 were among the top DEGs. There were differences in the number of DEGs identified per cancer type with the largest in bladder (n=4,938) and lowest in prostate (n=177) (Supplementary Table 1, 2). Pan-cancer DEGs had similar expression patterns across all malignancies (Figure 1a). Enrichment analysis (all DEGs) revealed that eight of the top-10 enriched pathways were ECM-related (Figure 1b). Further analysis confirmed hypoxia associated with all ECM gene categories at a pan-cancer level: collagens (9.8% of DEGs), proteoglycans (4.3%), glycoproteins (21.9%), regulators (26.8%), secreted factors (26.2%), and ECM-affiliated proteins (11%) (Figure 1c). Findings were consistent across all malignancies, except prostate, suggesting hypoxia-induced ECM alterations are cancer-prevalent.

**Figure 1.**
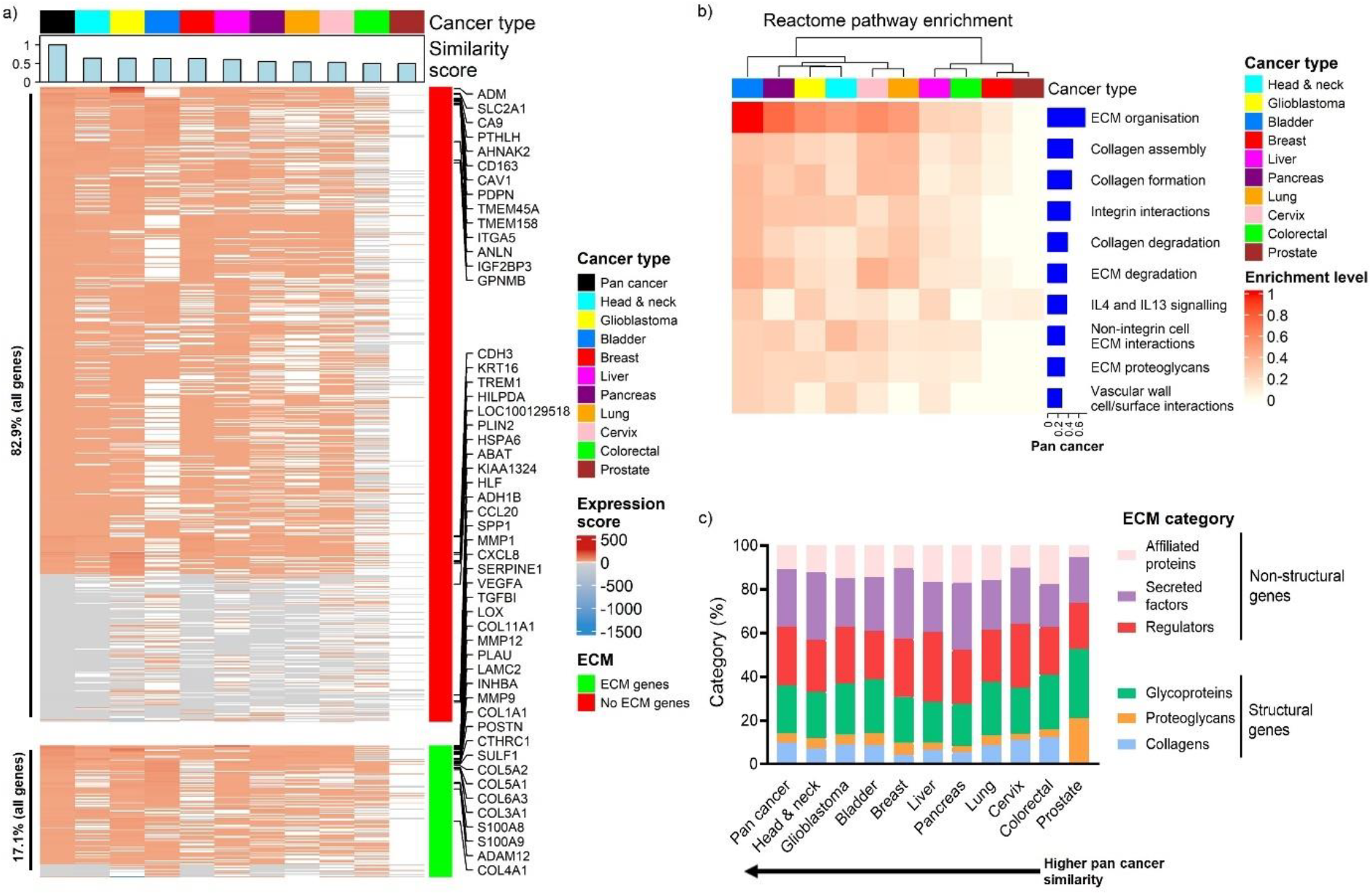
Meta-analysis of *in silico* datasets identifies prevalent extracellular matrix (ECM) pan-cancer alterations. (a) Transcriptome meta-analysis using random-effect modelling across 10 cancer types (n=131 cohorts; n=21,941 patients) identified 954 differentially expressed genes (DEGs) (fold change >1 or <-1; FDR<0.05; significance frequency >0.20) at a pan-cancer level. Top DEGs (fold change >2 or <-2; FDR<0.0001; significance frequency >0.3) gene symbols are highlighted. (b) Enrichment analysis highlighted ECM) pathways as commonly altered. (c) ECM ontology analysis reveals similar ECM alterations at transcriptomic level for all malignancies, except prostate.

### Hypoxia-induced ECM alterations *in vitro* recapitulate those found *ex vivo*

To further study hypoxia-induced ECM changes, we selected bladder cancer as model due to its high ECM pathway enrichment. *In vitro* approaches were used to explore ECM alterations under controlled conditions. Western blotting identified optimal conditions for *in vitro* work with seven days of chronic hypoxia (0.2% O_2_) inducing the greatest FN1 increase (Figure S1). Across all cell lines, hypoxia significantly affected 197 ECM genes at the RNA level and 127 at the protein level (p_adj_<0.05, FC >2 or FC<-2). Hierarchical (Figure S2) and k-means clustering (Figure S3) separated hypoxic and normoxic samples, highlighting hypoxia-associated ECM alterations. We identified 53 significant ECM genes (RNA and protein level), constituting our reference *in vitro* hypoxic-ECM atlas.

We crosschecked the atlas against bladder and pan-cancer *ex vivo* DEGs identified in Figure 1. Enrichment analysis showed hypoxia-associated changes *in vitro* recapitulated those *ex vivo* (Figure 2a, b). This similarity was dominated by HIF1/2-regulated ECM genes. More than 75% of *in vitro* genes were also identified *ex vivo*, with 23 genes found in all three groups (Figure 2c). Similar expression patterns were found for those 23 genes across all groups (Figure 2d). Interaction enrichment analysis identified key genes (*FN1, LAMC2, COL1A2, COL3A1, COL5A1, COL5A2*) linking “MET cell motility” and “MET/PTK2 signalling” with ECM-related pathways (Figure 2e), suggesting those ECM structural genes are common cancer regulators of cell motility during hypoxia. Altogether, results suggest our *in vitro* approach is a good model for studying hypoxia-associated ECM alterations.

**Figure 2.**
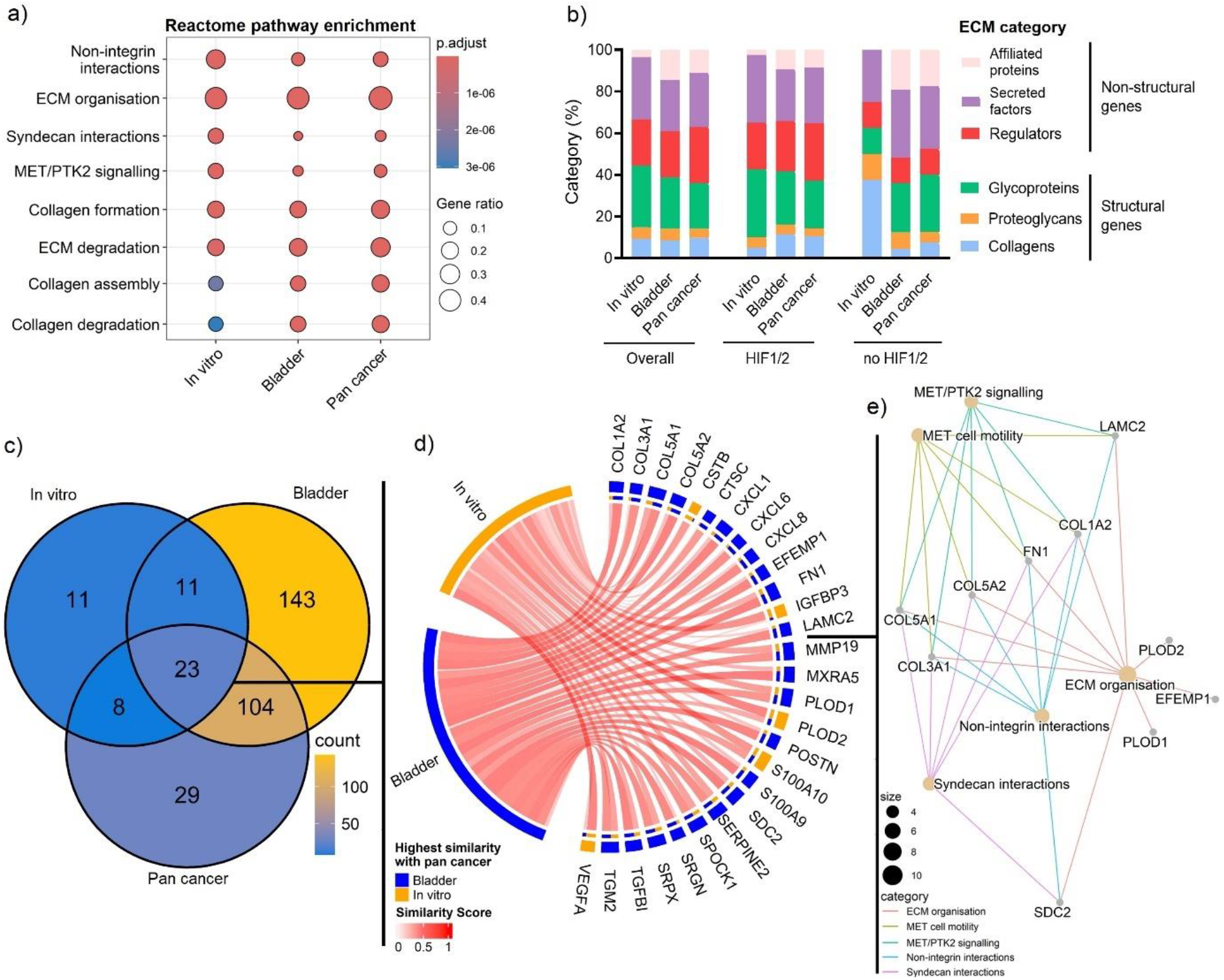
*In vitro* multi-omics recapitulate hypoxia-induced extracellular matrix (ECM) alterations found *ex vivo*. (a) Enrichment analysis shows hypoxia affects the same ECM-related pathways *in vitro* and *ex vivo*. (b) ECM ontology analysis shows that hypoxia affects similar ECM gene types in vitro and ex vivo, mainly driven by HIF1/2-regulated genes. (c) Venn diagram summarises overlap of hypoxia-associated genes. (d) Chord diagram shows similar expression patterns of the 23 genes identified *in vitro* and *ex vivo*. (e) Interaction enrichment analysis of the 23 common genes identified collagens (COL1A2, COL3A2, COL5A1, COL5A2), fibronectin (FN1) and laminin gamma-2 (LAMC2) as key proteins linking “ECM organisation” and “MET cell motility” pathways.

### Digital spatial profiling shows hypoxia affects gene expression differently in tumour and stroma

To independently confirm hypoxic-ECM changes in clinical samples whilst addressing stromal and tumour cells contributions, we analysed 79 ROIs (32 stroma, 47 core tumour) (Figure 3a). There were 260 DEGs (fold change ≥2 or ≤-2; p_adj_< 0.05) in CA9^+ve^ tumour, 142 DEGs in CA9^-ve^ tumour, and 27 DEGs in CA9^+ve^ stromal areas. We found no DEGs in CA9^-ve^ stroma (Figure 3b).

**Figure 3.**
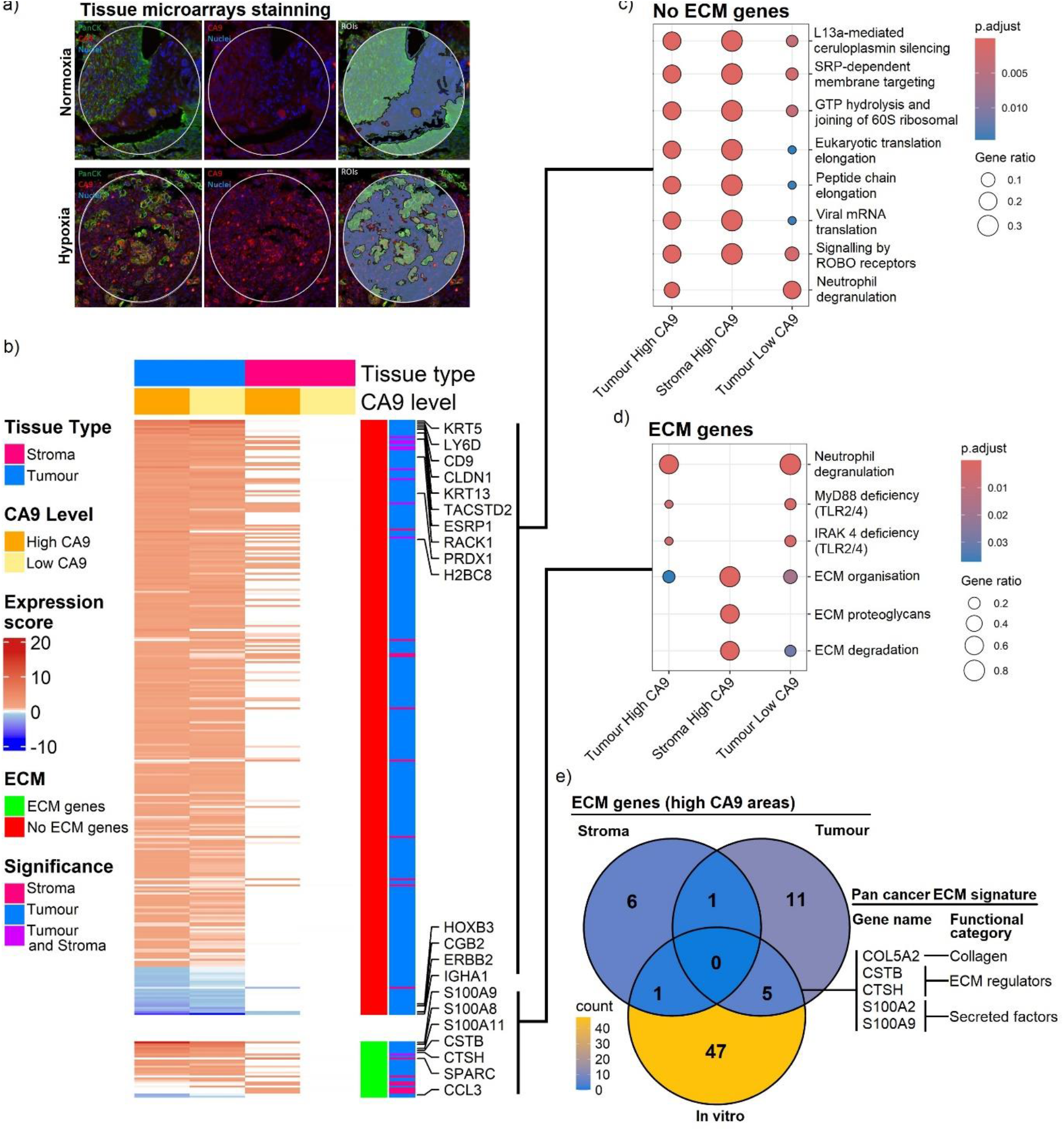
Digital spatial profiling shows hypoxia differently affects ECM gene expression in tumour and stroma. (a) Tissue microarrays including 79 ROIs (n=47 tumour, n=32 stroma) from 28 patients were stained for nuclei (blue), CA9 (red) to identify hypoxic areas and cytokeratin (green) to identify tumour cell enriched areas. (b) Spatial transcriptomics identified 260 differentially expressed genes (DEGs; fold change ≥2 or ≤-2; p_adj_< 0.05) in CA9^ve+^ tumour, 27 DEGs in CA9^ve+^ stromal, and 142 DEGs in CA9^ve-^ tumour areas. No DEGs were found in CA9^ve-^ stroma. Highly significant genes (p_adj_< 0.01, fold change ≥2 or ≤-2) are highlighted. (c) Enrichment analysis shows almost complete pathway overlapping in non-ECM DEGs for tumour and stromal areas. (d) Hypoxia changes in ECM genes modulate immune-related pathways in tumour but not stromal areas. (d) Only one gene (*ANXA2*) was significant in both CA9^ve+^ tumour and stromal areas. Five of the 53 genes from the *in vitro* hypoxic-ECM atlas were identified as significant in CA9^ve+^ tumour areas and used to constitute a pan-cancer hypoxic-ECM signature.

Enrichment analysis showed almost perfect pathway overlapping for non-ECM genes in CA9^+ve^ tumour and stromal areas, associating hypoxia with immune, metabolic, and protein translation pathways (Figure 3c). Differences were found in ECM genes. Commonly affected pathways were “ECM organisation” and “ECM degradation”. However, neutrophils and TLR2/4 immune pathways were identified only in CA9^+ve^ tumour areas, suggesting distinct hypoxia-associated ECM remodelling processes (Figure 3d). Of note, “neutrophil degranulation” was enriched significantly in bladder, colorectal, glioblastoma, liver and pan-cancer *ex vivo*, whilst toll-like receptors (TLR) pathways were identified in bladder, glioblastoma and lung *ex vivo* (Supplementary Table 3). *In vitro*, a protein-protein interaction (PPI) analysis predicted strong synergies between immune and ECM proteins in hypoxia across all cell lines (confidence score >0.7), altogether suggesting hypoxic tumour ECMs modulate immunity (Figure S4).

Comparison with the *in vitro* atlas identified further ECM-level differences. Only *ANXA2* (upregulated) was significant in CA9^+ve^ tumour and stromal areas. The greatest overlap was between hypoxic tumour areas and the atlas, which shared five ECM genes. These five hypoxia-associated ECM genes (*COL5A2, CSTB, CTSH, S100A2, S100A9*) formed a pan-cancer signature (Figure 3e).

### Hypoxic-ECM signature scores are prognostic and predictive of treatment benefit

Initial testing was performed in the BCON cohort, as the signature was identified using BCON samples. Cox regression showed patients with medium score values had worse prognoses. Tertile stratification was validated as cut-off in the remaining bladder radiotherapy cohorts and chosen for all analyses (Figure S6).

Univariable and multivariable analyses (Supplementary Table 4, 5) confirmed medium score patients had poor OS, MFS and ILRDFS in radiotherapy but not surgical bladder cohorts, suggesting the signature is predictive. Meta-analysis (n=1,514) showed the signature was prognostic for OS (medium scores HR=1.23, 95%CI=1.03-1.47, p=0.019) independently of treatment, stage, age and gender. Predictive capacity was tested after excluding non-muscle-invasive patients (not eligible for radiotherapy or radical surgery). The signature was independently predictive; medium score patients benefited from surgery over radiotherapy (Figure 4a). Kaplan-Meier confirmed radiotherapy improved OS for low (Figure 4b) and high (Figure 4c) scores. We then constructed an ANN model which predicted 10-year OS using treatment and hypoxic-ECM group (AUC=0.73; Figure 4d). The model improved after adding T stage, age, and concurrent chemotherapy as co-variables (AUC=0.79; Figure 4g).

**Figure 4:**
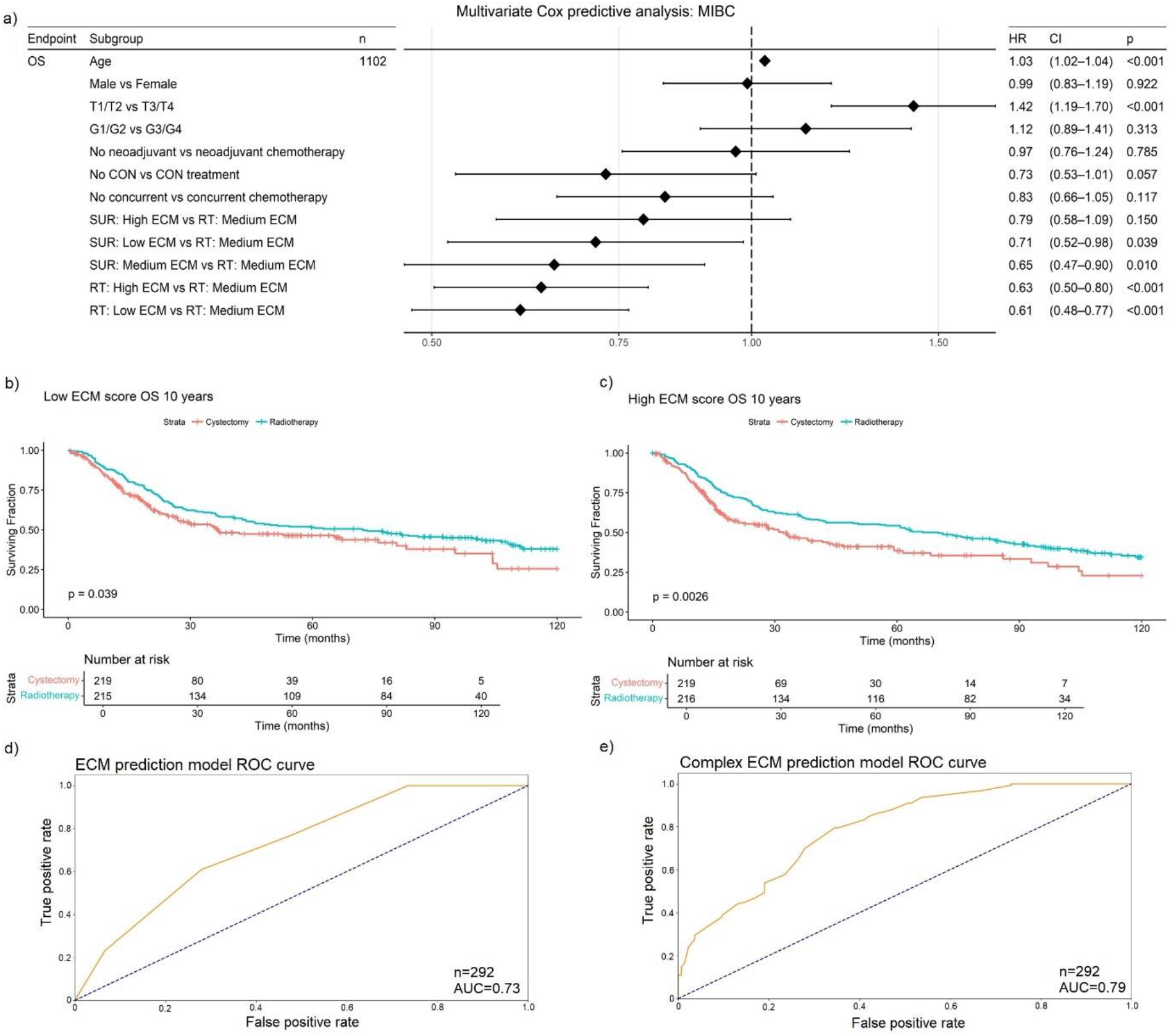
A candidate 5-gene hypoxic-ECM-associated signature predicts benefit from radiotherapy. A candidate signature (COL5A2, CSTB, CTSH, S100A2 and S100A9) was retrospectively validated in a combined MIBC cohort (n=1,259). Patients were tertile-stratified into high, medium and low groups based on the hypoxic-ECM signature median expression. (a) Multivariate Cox regression analysis showed a 35% decreased OS hazard ratio (HR) for medium score patients undergoing surgery instead of radiotherapy (p<0.010) independently of age, sex, stage, chemotherapy, and carbogen and nicotinamide (CON) treatment. (b, c) Kaplan-Meier analyses show better OS after radiotherapy for low (p=0.039) and high (p=0.0026) hypoxic-ECM scores, suggesting radiotherapy benefit. (f) A prediction model based on patient groups and treatment predicted 10-year OS (AUC=0.73). (g) Addition of T stage, age, and concurrent chemotherapy as variables increased the predictive capacity (AUC=0.79). The model was constructed using Artificial Neural Networks, with a random split of 80% for training, and 20% for validation.

As hypoxic-ECM alterations are common across cancers, we tested the signature in other malignancies. The signature was prognostic across multiple cohorts and cancer types in univariable and multivariable analyses (Supplementary Table 4, 5). Meta-analysis of the different cancers confirmed prognostic significance (glioblastoma, lung, pancreatic, prostate) with medium and high scores conferring a poor prognosis (Supplementary Table 6). Predictive capacity was explored for cancers with radiotherapy and surgery data. Low or high signature scores independently predicted radiotherapy benefit in breast (n=751; low scores HR=0.41, 95%CI=0.17-1.0, p=0.0498), head & neck (n=354; low scores HR=0.30, 95%CI=0.12-0.73, p=0.0085) and cervix (n=91; high scores HR=0.21, 95%CI=0.046-0.96, p=0.043) (Supplementary Table 6). Overall, our results suggest the pan-cancer signature is prognostic, with medium and high scores associating with worse outcomes, and predictive, with low and high scores associating with radiotherapy benefit.

### Hypoxic ECMs promote cell adhesion and impair cell migration, effects enhanced by radiation

As patients with high hypoxic-ECM scores had reduced metastasis and benefited from radiotherapy, we explored possible mechanisms *in vitro*. CDM coatings produced under severe hypoxia reduced cancer cell migration (Figure 5a-o) and increased cell adhesion (Figure 5p-s). Highest reduction in migration occurred in UMUC3, regardless of irradiation. For J82 and T24, migration impairment occurred only after irradiation and in a dose-dependent manner. RT4, a non-invasive cell line, was not assayed. The increase in cell adhesion was radiation-independent in all cell lines.

**Figure 5:**
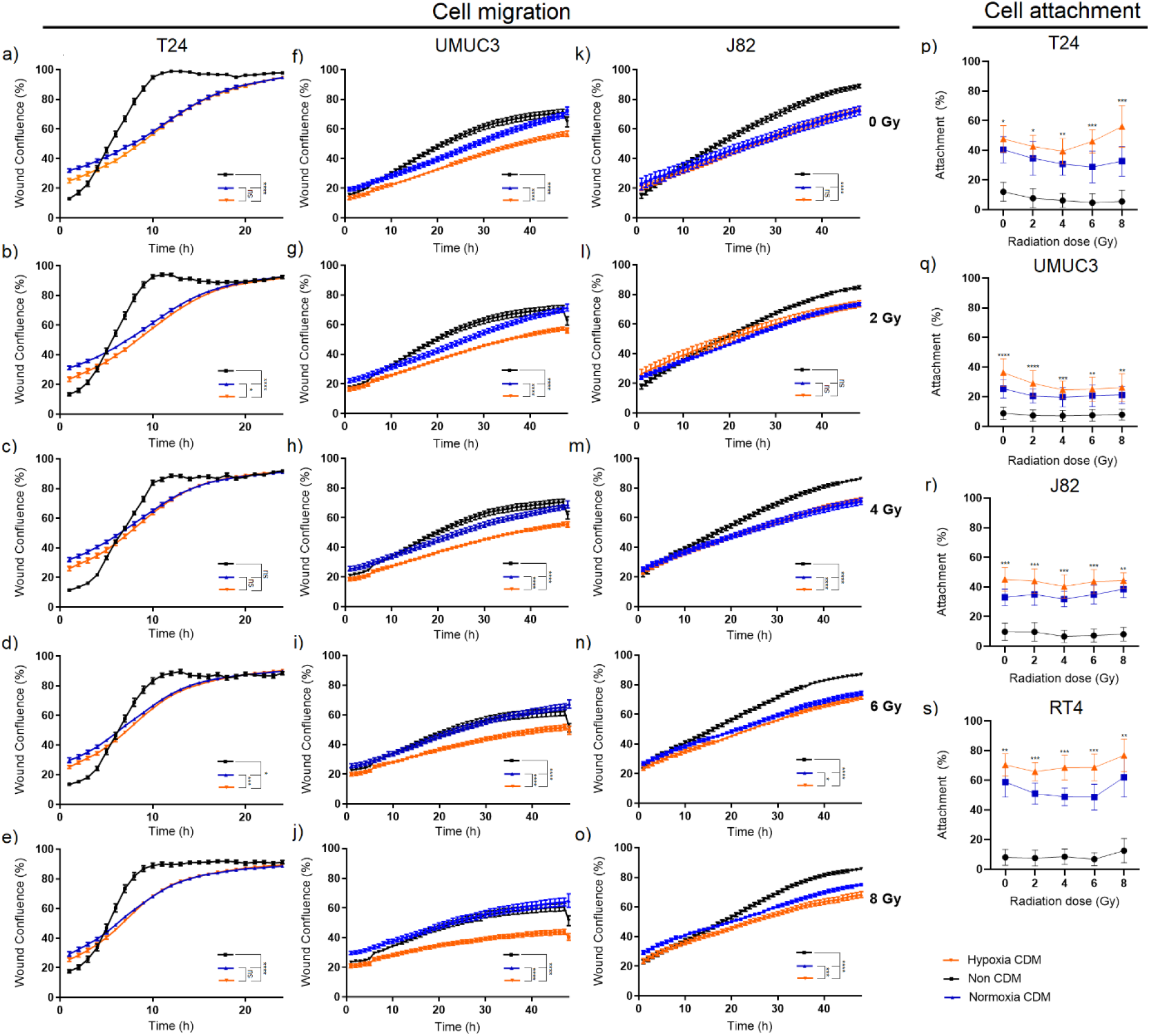
Hypoxic CDMs impair cell migration and promotes cell adhesion. Scratch assays for a-e) T24, f-j) UMUC3 and k-o) J82, and attachment assays for T24 (p), J82 (q), UMUC3 (r) and RT4 (s) cell lines compared their migration and adhesion capacities on hypoxic and normoxic CDM coatings, and a non-CDM coating control, for unirradiated and irradiated (0-8 Gy) cells. Graphs (f-j) show impaired cell migration for UMUC3 cells seeded onto hypoxic CDM (0-8 Gy) (p≤0.0001). Migration impairment was only significant after irradiation in T24 (2 Gy, p≤0.05; 6 Gy, p≤0.001) and J82 (6 Gy, p≤0.05; 8 Gy, p≤0.001). Significantly increased cell adhesion in hypoxic CDMs was found in all cell lines independently of radiation stress (p-s). Significance was assessed using 2-way ANOVA (scratch assays) and T-tests (attachment assays), with * for p≤0.05, ** for p≤0.01, *** for p≤0.001 and **** for p≤0.0001). Each data point represents mean±SEM of 3 independent experiments, with 12 (scratch assays) or 20 (attachment assays) technical replicates per experiment.

### Severe hypoxia reduces COL fibrogenesis, whilst radiation impairs cadherin, paxillin and vinculin co-localisation with ECM fibres

We associated COL5, COL1 and FN1 changes with “ECM organisation” and “MET cell motility” pathways. Hypoxia also affected collagen degradation assembly pathways. Hence, we investigated the effects of hypoxia on FN, COL5 and COL1 fibrogenesis and morphology (Figure S6). Hypoxia increased FN fibre number, width (T24, J82, RT4), and length (T24), while COL5 (T24, UMUC3) and COL1 (T24, J82, UMUC3) fibre numbers decreased, but increased in RT4. Fibre length increased for COL5 (UMUC3, RT4), whilst both length and width increased for COL1 (T24). FN, COL5, and COL1 levels increased during hypoxia in all cell lines.

To assess whether these changes affected FA signalling, we performed co-localisation analyses of E-cadherin, N-cadherin, and PXN/VCL in unirradiated (0 Gy) and irradiated (8 Gy) cells. Severely hypoxic ECMs affected the co-localisation area of: E-cadherin with COL5 (reduced in UMUC3, J82; increased in T24) and COL1 (reduced in UMUC3); N-cadherin with FN (reduced in T24; increased in RT4), COL1 (increased in UMUC3); PXN/VCL with FN (increased with T24), and COL5 (reduced in UMUC3). After irradiation, co-localisation was impaired. Radiation reduced co-localisation of: E-cadherin with FN (RT4), COL5 (all cell lines) and COL1 (T24, J82); N-cadherin with FN (RT4), COL5 and COL1 (UMUC3, RT4); and PXN/VCL with FN (T24) and COL5 (UMUC3). Similar results were found for signal intensity in co-localised areas. Interestingly, radiation increased E-cadherin co-localisation with FN in T24, and co-localised E-cadherin intensity with COL5 and COL1 in UMUC3 cells (Figures 6, S7, S8).

**Figure 6:**
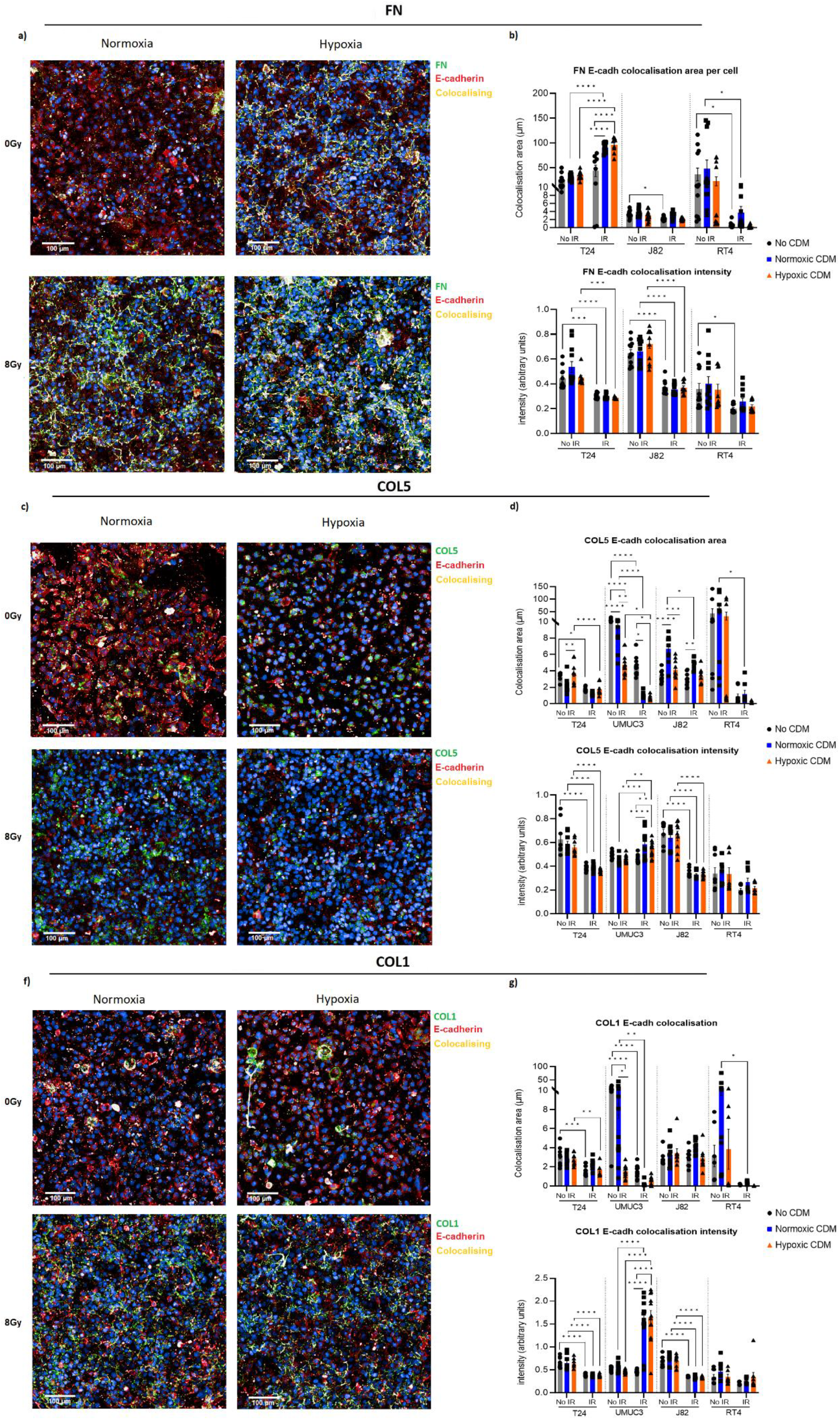
Hypoxic CDMs influence E-cadherin co-localisation with FN, COL5 and COL1 fibres, an effect impaired by irradiation. Figure shows immunofluorescence images for FN1 (a), COL5 (c) or COL1 (e) produced in hypoxia (0.2% O_2_) and normoxia (21% O_2_) and their co-localisation with E-cadherin in non-irradiated (0 Gy) and irradiated (8 Gy) T24 cells. Box plots show area and intensity of E-cadherin co-localised with FN1 (b), COL5 (d) and COL1 (g) fibres in T24, UMUC3, J82 and RT4 cells. Non-coated controls were included. No FN1 was detected for UMUC3. Hypoxic vs normoxic COL5 fibres had significantly increased (T24) or decreased (UMUC3, J82) co-localisation with E-cadherin in non-irradiated cells. Irradiation significantly decreased co-localised E-cadherin intensity in T24 and J82 for FN, COL5 and COL1 fibres. Increased co-localised E-cadherin area was seen for UMUC3 in COL5 and COL1 fibres. After irradiation E-cadherin co-localisation area decreased with FN1 (RT4), COL5 (T24, UMUC3, J82, RT4) and COL1 (T24, UMUC3, RT4). Radiation increased E-cadherin co-localisation in T24 with FN1. Significance is defined as p_adj_<0.05 and at least 20% fold-change, with * for p_adj_<0.05, ** for p_adj_<0.01, *** for p_adj_<0.001 and **** for p_adj_<0.0001. Signal intensity and colocalisation area were normalised to total cell number. Data are for three independent experiments, each with three technical repeats.

Radiation reduced total expression levels of E-cadherin (UMUC3, RT4) and PXN/VCL (UMUC3), but not N-cadherin (Figure S9). Our data suggest severe hypoxia promotes FN1, but impairs COL5 and COL1 fibrogenesis. Radiation altered cell/ECM interactions, reducing PXN/VCL/cadherin co-localisation with ECM fibres.

## Discussion

The major finding in this manuscript is that ECM changes are a dominant effect of hypoxia in cancer, *in silico* linking hypoxia-induced ECM alterations with immune infiltration and cell migration via MET signalling. In clinical samples, hypoxic tumour and stromal regions showed distinct ECM changes, associated with immune pathways only in tumour areas. Hypoxia-induced ECM changes directly impacted outcomes and treatment benefit; we identified a 5-gene hypoxic-ECM signature, prognostic and predictive of radiotherapy benefit across multiple cancers. Unexpectedly, patients with high hypoxic-ECM scores benefited from radiotherapy versus surgery and had reduced metastasis, a concept validated *in vitro* and related to radiation impairment of cell/ECM interactions.

Although hypoxia’s role in ECM remodelling is well-documented and related to angiogenesis^1,45,46^, a pan-cancer analysis has not been performed. Our study identified hypoxia-driven commonalities, providing a reference atlas. Glycoproteins (e.g. FN1, SDCs) were the most affected ECM genes. While prior hypoxia studies mostly focused on collagens as drivers of fibrosis and metastasis^1,10,45^, our data suggest glycoproteins may be more important. This concept supports recent findings showing HIF1 upregulates the glycoprotein SDC3, correlating its expression with hypoxia-scores and a poor prognosis^47^.

ECM regulators and secreted factors comprised 25% of hypoxia-regulated ECM genes, aligning with reports of hypoxia-induced ECM remodelling promoting their release^1,10,45^. HIF1/2 regulated over 70% of hypoxia-associated ECM genes across all ECM families, supporting their reported role in ECM regulation^1,10,45^. A novel finding was that HIF1 and HIF2 predominantly targeted the same ECM genes, suggesting equal importance for ECM regulation. Notably, 25% of ECM genes were HIF1/2-independent, implying other mechanisms, e.g. UPR, are also key ECM regulators. For instance, several COLs (e.g., COL1A2, COL3A1, COL4A5) were not directly regulated by HIF1/2, consistent with studies linking HIF3 and UPR signalling with increased COL and FN production^48,49^.

*In vitro* and *ex vivo* patient sample data strongly overlapped, with more hypoxia-regulated ECM genes in tumour vs stromal regions. Since central tumour areas are more likely to be hypoxic^50^, cancer cells may play a dominant role in hypoxia-induced ECM changes. Furthermore, in prostate cancer, there are differences in the composition of the ECM produced by tumour cells and fibroblasts^51^. Our data identified ECM compositional differences, with hypoxia influencing immune pathways only in tumour areas. A consensus subset of hypoxia-regulated ECM genes linked with MET signalling, a modulator of immune cell migration^52^. Indeed, hypoxia enhances neutrophil and macrophage recruitment to promote ECM remodelling and angiogenesis^1,45^, which associates with immunosuppression and a poor prognosis in bladder cancer^53^. Although exact mechanisms are unclear, our data highlights MET signalling as a mediator pathway. We predicted interactions among cytokines (e.g. CXCL8) and structural ECM proteins (e.g. FN1, COL5A2) *in vitro*, which all increased under hypoxia. Jing *et al*. showed CXCL8 induces neutrophil recruitment and polarisation^54^. Therefore, MET and chemokine signalling may have synergies regulating myeloid cell recruitment and immunosuppression, which could be targeted.

Our hypoxic-ECM signature was independently prognostic and predictive of radiotherapy benefit in multiple cancers. Only the radiosensitivity index predicts radiotherapy benefit across several cancers^55^. Additionally, there are no validated radiosensitivity signatures for bladder^55^. High hypoxic-ECM scores predicted radiotherapy benefit, contradicting the known link between hypoxia and radioresistance. Although counter-intuitive, Zolzer and Streffer showed severe hypoxia increases radiosensitivity *in vitro* if irradiation is not performed under hypoxia^56^.

To understand these findings, we evaluated the capacity of severely hypoxic ECMs to modulate cell migration after irradiation, showing that severely hypoxic ECMs decreased cell migration and increased cell adhesion. Although increased FN1 and COL deposition may be sufficient to promote cell attachment, COL fibrogenesis and crosslinking are required for cancer cells to migrate and invade healthy tissues^57^. Indeed, hypoxia promoted cell migration through MET signalling in three-dimensional collagen-branched hydrogel^58^. Hence, we investigated changes in COL and FN1 fibrogenesis. We found hypoxia impaired COL fibrogenesis, supporting previous studies in several cancers (breast^16,17^, prostate^17^, vulval^18^, head & neck^18^), some showing decreased cell migration^18^. Although hypoxia increased COL and FN1 fibre length and width, the fibres were immature (length ∼13 μm)^59^. We also showed irradiation-impaired PXN/VCL/cadherin co-localisation with ECM fibres in hypoxic ECMs. Cadherins, VCL and PXN spatial location regulate FA formation^11^, implying that reduced co-localisation with ECM fibres impairs FAs formation. Other studies showed radiotherapy-induced changes in FAs depend on the ECM composition^60,61^, with low stiffness increasing adhesion and decreasing migration of irradiated cancer cells^61^. As COL fibrogenesis correlates with matrix stiffness^10,12^, the observed impairment would decrease stiffness, reducing cell migration after irradiation. Overall, our data suggest that severe hypoxia reduces ECM stiffness, while radiation weakens cell/ECM interactions, impairing metastasis. These results explain the reduced metastasis in high hypoxic-ECM score patients, though medium-score patients had increased risk. COL crosslinking and fibrogenesis might occur within specific oxygen ranges, a hypothesis warranting further investigation. Furthermore, FA signalling of irradiated cells on hypoxic-ECMs is largely unexplored, highlighting the need for mechanistic studies to identify targets for hypoxic-ECM-directed therapies.

Of note, UMUC3, a non-FN1-producing cell line, showed the highest impairment of cell migration. ITGA5, a key FN1 receptor, was among the top upregulated pan-cancer genes under hypoxia. We also observed increased FN1 fibrogenesis and PXN/VCL colocalisation with FN1 *in vitro*. Increased FN1 and ITGA5 expression in hypoxia have been respectively linked to increased cell migration^62^ and breast cancer metastasis^63^. ITGA5/FN1 signalling may regulate cancer cell migration under severe hypoxia when COL fibres are lacking. Functional experiments (e.g. *in vitro/in vivo* ITGA5 blockade) are needed to confirm this hypothesis.

Our study has limitations. The transcriptomic meta-analysis relied on 10 hypoxia signatures, assuming equal stratification capacity. Nevertheless, well-known hypoxia markers (e.g., CA9, SLC2A1)^64,65^ ranked among the top upregulated genes, validating our approach. Although our ChIP-seq was limited to one cell line (T24), evidence shows HIF1/2 binding to DNA regions is highly conserved^66^. ChIP-seq focused on proximal promoter regions. However, HIF1 also binds distant regulatory regions^67,68^, following dynamic temporal binding fluctuations^48,69^. Future studies should explore non-HIF1/2 mechanisms and HIF1/2 distal binding and temporal dynamics. The links between hypoxia, ECM remodelling, and immunity also require functional validation, as they were predicted *in silico*.

## Conclusion

Hypoxia induces prevalent pan-cancer ECM alterations driven by tumour cells. These changes have clinical relevance and modulate radiotherapy benefit, as shown by a newly identified pan-cancer hypoxic-ECM signature. Highly hypoxic ECMs associated with radiotherapy benefit and reduced metastasis, a concept validated *in vitro* and associated with impaired COL fibrogenesis and FA formation. Prospective validation of the signature is warranted, alongside mechanistic studies to therapeutically exploit the synergies between hypoxic ECMs and radiotherapy.

## Supporting information

Supplementary results, materials and methods

## Acknowledgements

The work was funded by the NIHR Manchester Biomedical Research Centre (NIHR129943). The work was also supported by Cancer Research UK Major Centre (C147/A25254) and project grant (C1098/A9437; C2094/A11365; C13329/A21671) funding. The support of D. Knight, E. Keevill and J.N. Selley at the Bio-MS mass spectrometry core facility (RRID: SCR_020987) in the Faculty of Biology, Medicine and Health at the University of Manchester is gratefully acknowledged. The mass spectrometers and microscopes used in this study were purchased with grants from BBSRC, Wellcome Trust and the University of Manchester Strategic Fund.

Work was carried out at The University of Manchester, Cancer Research UK – Manchester Institute, and the Manchester Cancer Research Centre, which provided infrastructure support and access to core facilities for the experiments performed in this study.

## Author Contributions

Conceptualisation: C.G.Q., J.G.A., V.B., J.D.H., M.J.H., A.C., C.M.W.; Methodology: C.G.Q., J.G.A., A.O., V.B., A.S.F., J.D.H., M.J.H., A.C., C.M.W.; Software: C.G.Q., A.S.F, M.R.; Validation: C.G.Q., A.S.F.; Formal analysis: C.G.Q., J.G.A., A.O., A.S.F., M.R., A.B.; Investigation: C.G.Q., J.G.A., A.O., V.S., R.S., A.B., M.E., G.M.; Resources: T.L., P.H., N.D.J., E.H., R.A.H., N.P., A.C.; Data curation: C.G.Q., T.L., M.R.; Visualisation: C.G.Q., A.B.; Supervision: T.A.D.S., V.B., J.D.H., P.H., M.J.H., A.C., C.M.W.; Project administration: K.R., J.D.H., M.J.H., A.C., C.M.W.; Funding acquisition: K.R., J.D.H., M.J.H., A.C., C.M.W.; Writing (original draft): C.G.Q.; Writing (review & editing): M.R., A.B., V.B., T.A.D.S., J.D.H., M.J.H., A.C., C.M.W.

## Statement of data availability

*In vitro* proteomic data will be made publicly available in the ProteomeXchange repository. *In vitro* and Christie-GemX transcriptomic data will be made available in the GEO repository. BCON, BC2001, and Christie-GemX clinical data are available upon request from the clinical leads of these studies. All remaining datasets and outcome data are available in public repositories.

## Statement of code availability

The code used in the analyses of this study will be made available in the GitHub repository.

## References

1. Bigos, K. J. A. et al. Tumour response to hypoxia: understanding the hypoxic tumour microenvironment to improve treatment outcome in solid tumours. Front Oncol 14, 1331355 (2024).

2. Gray, L.H., Conger, A.D., Ebert, M., Hornsey, S. & Scott, O. C. The Concentration of Oxygen Dissolved in Tissues at the Time of Irradiation as a Factor in Radiotherapy. http://dx.doi.org/10.1259/0007-1285-26-312-638 26, 638–648 (1953).

3. Boulefour, W. et al. A Review of the Role of Hypoxia in Radioresistance in Cancer Therapy. Med Sci Monit 27, (2021).

4. Keith, B., Johnson, R. S. & Simon, M. C. HIF1α and HIF2α: sibling rivalry in hypoxic tumour growth and progression. Nat Rev Cancer 12, 9–22 (2011).

5. Bolland, H., Ma, T. S., Ramlee, S., Ramadan, K. & Hammond, E. M. Links between the unfolded protein response and the DNA damage response in hypoxia: a systematic review. Biochem Soc Trans 49, 1251–1263 (2021).

6. Denko, N. C. et al. Investigating hypoxic tumor physiology through gene expression patterns. Oncogene 22, 5907–5914 (2003).

7. Shi, R., Liao, C. & Zhang, Q. Hypoxia-Driven Effects in Cancer: Characterization, Mechanisms, and Therapeutic Implications. Cells 10, 1–26 (2021).

8. He, Y. et al. Tumor-Associated Extracellular Matrix: How to Be a Potential Aide to Anti-tumor Immunotherapy? Front Cell Dev Biol 9, (2021).

9. Robertson, J. et al. Defining the phospho-adhesome through the phosphoproteomic analysis of integrin signalling. Nat Commun 6, 1–13 (2015).

10. Gilkes, D. M., Wirtz, D. & Semenza, G. L. Hypoxia and the extracellular matrix: drivers of tumour metastasis. Nat Rev Cancer 14, 430–439 (2015).

11. Mui, K. L., Chen, C. S. & Assoian, R. K. The mechanical regulation of integrin–cadherin crosstalk organizes cells, signaling and forces. J Cell Sci 129, 1093 (2016).

12. Pickup, M. W., Mouw, J. K. & Weaver, V. M. The extracellular matrix modulates the hallmarks of cancer. EMBO Rep 15, 1243–1253 (2014).

13. Dickreuter, E. et al. Targeting of β1 integrins impairs DNA repair for radiosensitization of head and neck cancer cells. Oncogene 2016 35:11 35, 1353–1362 (2015).

14. Storch, K. & Cordes, N. Focal adhesion-chromatin linkage controls tumor cell resistance to radio- and chemotherapy. Chemother Res Pract 2012, 1–10 (2012).

15. Qiao, L. et al. Targeting Epithelial-to-Mesenchymal Transition in Radioresistance: Crosslinked Mechanisms and Strategies. Front Oncol 12, (2022).

16. Goggins, E. et al. Hypoxia Inducible Factors Modify Collagen I Fibers in MDA-MB-231 Triple Negative Breast Cancer Xenografts. Neoplasia 20, 131–139 (2018).

17. Kakkad, S. M. et al. Hypoxic tumor microenvironments reduce collagen I fiber density. Neoplasia 12, 608–617 (2010).

18. Madsen, C. D. et al. Hypoxia and loss of PHD2 inactivate stromal fibroblasts to decrease tumour stiffness and metastasis. EMBO Rep 16, 1394–1408 (2015).

19. Yang, L. et al. A Gene Signature for Selecting Benefit from Hypoxia Modification of Radiotherapy for High-Risk Bladder Cancer Patients. Clinical Cancer Research 23, 4761–4768 (2017).

20. Smith, T. A. D. et al. A hypoxia biomarker does not predict benefit from giving chemotherapy with radiotherapy in the BC2001 randomised controlled trial. EBioMedicine 101, (2024).

21. Buffa, F. M., Harris, A. L., West, C. M. & Miller, C. J. Large meta-analysis of multiple cancers reveals a common, compact and highly prognostic hypoxia metagene. Br J Cancer 102, 428– 435 (2010).

22. Yang, Y., Li, Y., Qi, R. & Zhang, L. Constructe a novel 5 hypoxia genes signature for cervical cancer. Cancer Cell Int 21, (2021).

23. Chen, P. et al. Identification of Hypoxia-Associated Signature in Colon Cancer to Assess Tumor Immune Microenvironment and Predict Prognosis Based on 14 Hypoxia-Associated Genes. Int J Gen Med 16, 2503–2518 (2023).

24. Tardón, M. C. et al. An Experimentally Defined Hypoxia Gene Signature in Glioblastoma and Its Modulation by Metformin. Biology (Basel) 9, 1–17 (2020).

25. Chang, W. H., Forde, D. & Lai, A. G. A novel signature derived from immunoregulatory and hypoxia genes predicts prognosis in liver and five other cancers. J Transl Med 17, (2019).

26. Eustace, A. et al. A 26-gene hypoxia signature predicts benefit from hypoxia-modifying therapy in laryngeal cancer but not bladder cancer. Clin Cancer Res 19, 4879–4888 (2013).

27. Lane, B., Khan, M. T., Choudhury, A., Salem, A. & West, C. M. L. Development and validation of a hypoxia-associated signature for lung adenocarcinoma. Sci Rep 12, (2022).

28. Abou Khouzam, R. et al. An Eight-Gene Hypoxia Signature Predicts Survival in Pancreatic Cancer and Is Associated With an Immunosuppressed Tumor Microenvironment. Front Immunol 12, 1861 (2021).

29. Yang, L. et al. Development and Validation of a 28-gene Hypoxia-related Prognostic Signature for Localized Prostate Cancer. EBioMedicine 31, 182–189 (2018).

30. Forker, L. J. et al. Technical development and validation of a clinically applicable microenvironment classifier as a biomarker of tumour hypoxia for soft tissue sarcoma. Br J Cancer 128, 2307–2317 (2023).

31. Love, M. I., Huber, W. & Anders, S. Moderated estimation of fold change and dispersion for RNA-seq data with DESeq2. Genome Biol 15, 1–21 (2014).

32. Ritchie, M. E. et al. limma powers differential expression analyses for RNA-sequencing and microarray studies. Nucleic Acids Res 43, e47–e47 (2015).

33. Storey, J. D. & Tibshirani, R. Statistical significance for genomewide studies. Proc Natl Acad Sci U S A 100, 9440–9445 (2003).

34. Viechtbauer, W. Conducting Meta-Analyses in R with the metafor Package. J Stat Softw 36, 1– 48 (2010).

35. Shao, X. et al. MatrisomeDB 2.0: 2023 updates to the ECM-protein knowledge database. Nucleic Acids Res 51, D1519–D1530 (2023).

36. Robertson, J. et al. Defining the phospho-adhesome through the phosphoproteomic analysis of integrin signalling. Nat Commun 6, (2015).

37. Yang, L. et al. Development and Validation of a 28-gene Hypoxia-related Prognostic Signature for Localized Prostate Cancer. EBioMedicine 31, 182–189 (2018).

38. Humphries, J. D. et al. Pancreatic ductal adenocarcinoma cells employ integrin α6β4 to form hemidesmosomes and regulate cell proliferation. Matrix Biol 110, 16–39 (2022).

39. Smith, V. et al. Hypoxia Is Associated with Increased Immune Infiltrates and Both Anti-Tumour and Immune Suppressive Signalling in Muscle-Invasive Bladder Cancer. Int J Mol Sci 24, (2023).

40. Byron, A. et al. A proteomic approach reveals integrin activation state-dependent control of microtubule cortical targeting. Nat Commun 6, (2015).

41. Hoskin, P. J., Rojas, A. M., Bentzen, S. M. & Saunders, M. I. Radiotherapy with concurrent carbogen and nicotinamide in bladder carcinoma. J Clin Oncol 28, 4912–4918 (2010).

42. Eustace, A. et al. Necrosis predicts benefit from hypoxia-modifying therapy in patients with high risk bladder cancer enrolled in a phase III randomised trial. Radiother Oncol 108, 40–47 (2013).

43. Hunter, B. A. et al. Expression of hypoxia-inducible factor-1α predicts benefit from hypoxia modification in invasive bladder cancer. Br J Cancer 111, 437–443 (2014).

44. Eyers, M. et al. Digital spatial profiling of the microenvironment of muscle invasive bladder cancer. Commun Biol 7, (2024).

45. Winkler, J., Abisoye-Ogunniyan, A., Metcalf, K. J. & Werb, Z. Concepts of extracellular matrix remodelling in tumour progression and metastasis. Nature Communications 2020 11:1 11, 1– 19 (2020).

46. Pugh, C. W. & Ratcliffe, P. J. Regulation of angiogenesis by hypoxia: role of the HIF system. Nat Med 9, 677–684 (2003).

47. Prieto-Fernández, E. et al. Hypoxia Promotes Syndecan-3 Expression in the Tumor Microenvironment. Front Immunol 11, (2020).

48. Wiafe, B. et al. Hypoxia-increased expression of genes involved in inflammation, dedifferentiation, pro-fibrosis, and extracellular matrix remodeling of human bladder smooth muscle cells. In Vitro Cellular & Developmental Biology - Animal 2016 53:1 53, 58–66 (2016).

49. Maiers, J. L. et al. The unfolded protein response mediates fibrogenesis and collagen I secretion through regulating TANGO1 in mice. Hepatology 65, 983–998 (2017).

50. Saxena, K. & Jolly, M. K. Acute vs. Chronic vs. cyclic hypoxia: Their differential dynamics, molecular mechanisms, and effects on tumor progression. Biomolecules 9, (2019).

51. Tian, C. et al. Proteomic analyses of ECM during pancreatic ductal adenocarcinoma progression reveal different contributions by tumor and stromal cells. Proc Natl Acad Sci U S A 116, 19609–19618 (2019).

52. Hamouda, A. E. I. et al. Met-Signaling Controls Dendritic Cell Migration in Skin by Regulating Podosome Formation and Function. Journal of Investigative Dermatology 143, 1548-1558.e13 (2023).

53. Smith, V. et al. Hypoxia Is Associated with Increased Immune Infiltrates and Both Anti-Tumour and Immune Suppressive Signalling in Muscle-Invasive Bladder Cancer. Int J Mol Sci 24, (2023).

54. Jing, W. et al. Tumor-neutrophil cross talk orchestrates the tumor microenvironment to determine the bladder cancer progression. Proc Natl Acad Sci U S A 121, e2312855121 (2024).

55. Bleaney, C. W. et al. Clinical Biomarkers of Tumour Radiosensitivity and Predicting Benefit from Radiotherapy: A Systematic Review. Cancers 2024, Vol. 16, Page 1942 16, 1942 (2024).

56. Zölzer, F. & Streffer, C. Increased radiosensitivity with chronic hypoxia in four human tumor cell lines. Int J Radiat Oncol Biol Phys 54, 910–920 (2002).

57. Song, K. et al. Collagen Remodeling along Cancer Progression Providing a Novel Opportunity for Cancer Diagnosis and Treatment. Int J Mol Sci 23, (2022).

58. Pennacchietti, S. et al. Hypoxia promotes invasive growth by transcriptional activation of the met protooncogene. Cancer Cell 3, 347–361 (2003).

59. Kadler, K. E., Holmes, D. F., Trotter, J. A. & Chapman, J. A. Collagen fibril formation. Biochemical Journal 316, 1 (1996).

60. Cordes, N., Hansmeier, B., Beinke, C., Meineke, V. & Van Beuningen, D. Irradiation differentially affects substratum-dependent survival, adhesion, and invasion of glioblastoma cell lines. Br J Cancer 89, 2122–2132 (2003).

61. Panzetta, V. et al. Adhesion and Migration Response to Radiation Therapy of Mammary Epithelial and Adenocarcinoma Cells Interacting with Different Stiffness Substrates. Cancers (Basel) 12, (2020).

62. Lin, T. C. et al. Fibronectin in Cancer: Friend or Foe. Cells 9, (2020).

63. Ju, J. A. et al. Hypoxia Selectively Enhances Integrin α5β1 receptor expression in breast cancer to promote metastasis. Molecular Cancer Research 15, 723–734 (2017).

64. Le, Q. T. & Courter, D. Clinical biomarkers for hypoxia targeting. Cancer and Metastasis Reviews 27, 351–362 (2008).

65. Kim, J. Il et al. Expression of hypoxic markers and their prognostic significance in soft tissue sarcoma. Oncol Lett 9, 1699–1706 (2015).

66. Smythies, J. A. et al. Inherent DNA-binding specificities of the HIF-1α and HIF-2α Transcription factors in chromatin. EMBO Rep 20, (2019).

67. Platt, J. L. et al. Capture-C reveals preformed chromatin interactions between HIF-binding sites and distant promoters. EMBO Rep 17, 1410–1421 (2016).

68. Andrysik, Z., Bender, H., Galbraith, M. D. & Espinosa, J. M. Multi-omics analysis reveals contextual tumor suppressive and oncogenic gene modules within the acute hypoxic response. Nature Communications 2021 12:1 12, 1–18 (2021).

69. Jeknić, S., Kudo, T., Song, J. J. & Covert, M. W. An optimized reporter of the transcription factor hypoxia-inducible factor 1α reveals complex HIF-1α activation dynamics in single cells. J Biol Chem 299, 104599 (2023).

70. Humphries, J. D. et al. Pancreatic ductal adenocarcinoma cells employ integrin α6β4 to form hemidesmosomes and regulate cell proliferation. Matrix Biology 110, 16–39 (2022).

71. Robertson, J. et al. Characterization of the Phospho-Adhesome by Mass Spectrometry-Based Proteomics. Methods Mol Biol 1636, 235–251 (2017).

72. Byron, A. et al. A proteomic approach reveals integrin activation state-dependent control of microtubule cortical targeting. Nat Commun 6, 1–14 (2015).

73. Leek JT et al. sva: Surrogate Variable Analysis v.3.50.0. Bioconductor (2023) doi:10.18129/B9.bioc.sva.

74. Law, C. W., Chen, Y., Shi, W. & Smyth, G. K. voom: Precision weights unlock linear model analysis tools for RNA-seq read counts. Genome Biol 15, (2014).

75. Shannon, P. et al. Cytoscape: a software environment for integrated models of biomolecular interaction networks. Genome Res 13, 2498–2504 (2003).

76. Franceschini, A. et al. STRING v9.1: protein-protein interaction networks, with increased coverage and integration. Nucleic Acids Res 41, (2013).

77. Pezoulas, V. C. et al. A computational pipeline for data augmentation towards the improvement of disease classification and risk stratification models: A case study in two clinical domains. Comput Biol Med 134, 104520 (2021).

