## Supplementary results, materials and methods for "Hypoxia-induced extracellular matrix changes are conserved in cancer and directly impact radiotherapy benefit"

**f)** Medicines Discovery Catapult, Alderley Park, Cheshire, UK.

**g)** Nuclear Futures Institute, Bangor University, Bangor LL57 2DG, UK.

**h)** The Institute of Cancer Research, London SW3 6JJ, UK.

**i)** Department of Life Sciences, Manchester Metropolitan University, Manchester M1 5GD, UK.

*Authors equally contributed to the production of the manuscript.

### Supplementary figures

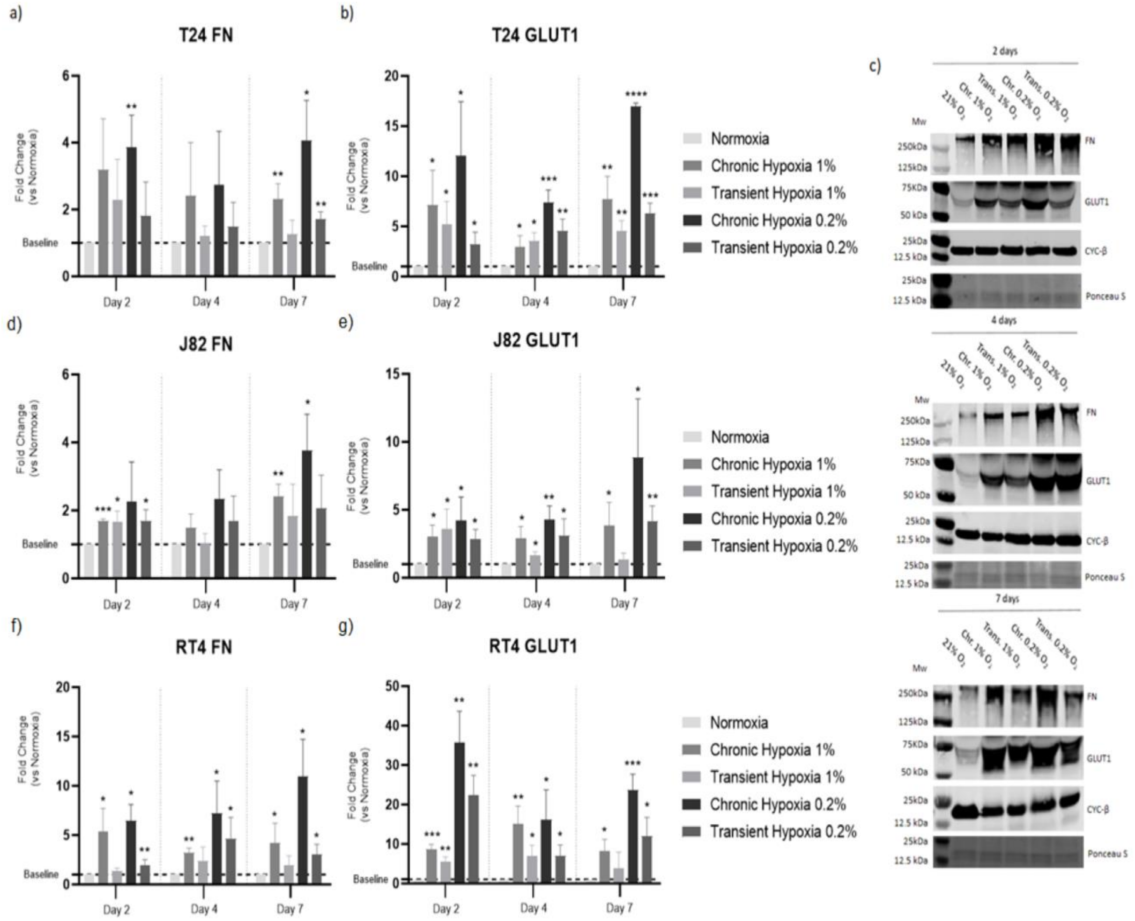


**Figure S1: Hypoxia-induced changes in fibronectin (FN) and glucose transporter 1 (GLUT1) protein expression in bladder cancer cells.** Cells were grown for two, four and seven days in chronic hypoxia (1% and 0.2% $O_2$), transient hypoxia (1% and 0.2% $O_2$) or normoxia (20.9% $O_2$). Fold changes in FN protein increased in hypoxia (a, d, f) and were mirrored by GLUT1 fold change increases (b, e, f). Western blot images also provided visual evidence of FN and GLUT1 increases in hypoxia (c). Highest FN fold change was seen for chronic hypoxia at 0.2 % $O_2$, with the maximum fold change occurring at day seven (a, d, f). The highest GLUT1 fold change was also seen for chronic hypoxia at 0.2 % $O_2$, with the maximum fold change also occurring at day 7 for T24 (b) and J82 (e), but at day 2 for RT4 (g). Western blots images (c) are representative images from the T24 cell line. Cyclophilin β (CYC-β) was used for normalisation as a loading control for GLUT1. FN levels were normalised through Ponceau S staining total protein quantification. Fold changes were calculated using normoxia signal intensity levels as reference. Each bar represents the average and the standard deviation for three biological repeats. Shapiro-Wilks tests were used to address data normality. Unpaired t tests were used to determine statistical significance (* for $p \le 0.05$, ** for $p \le 0.01$, *** for $p \le 0.001$, **** for $p \le 0.0001$).

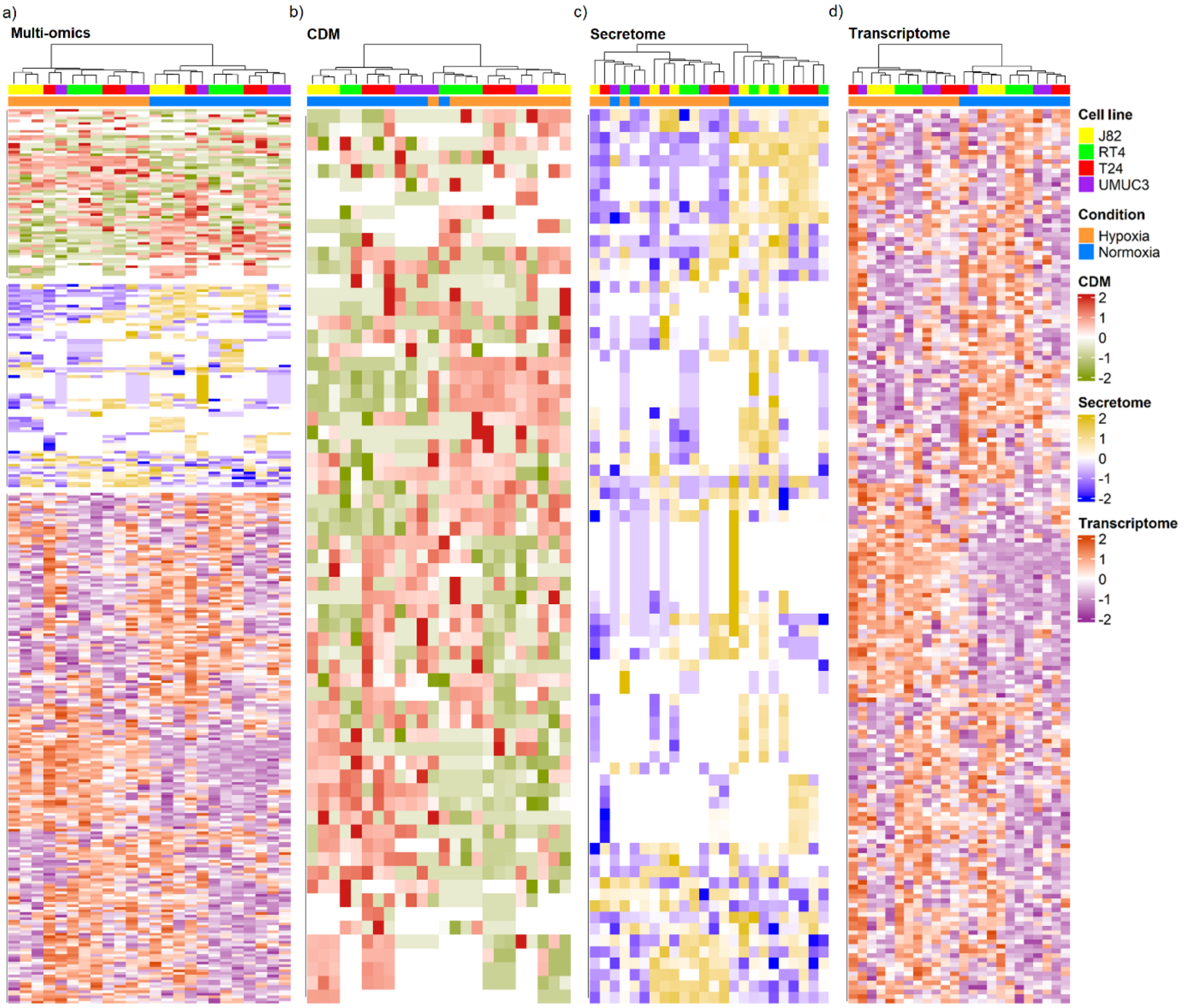


**Figure S2: Hypoxia modulates extracellular matrix (ECM) composition through changes at the RNA and protein level.** Unsupervised Z-scores clustering of multi-omics (a), cell-derived ECM (CDM) (b), secretome (c), and transcriptome (d) datasets from significant ($p_{adj} \leq 0.05$, fold change ≥2 or ≤-2) RNAs (n=196), and proteins (n=121, 66 CDM, 67 secretome) identified in at least in one cell line. RNAs were detected using whole-cell transcriptomics and proteins by proteomic analysis of CDMs and secretomes. T24, UMUC3, RT4 and J82 cells were cultured for 24 h (transcriptomics) or 7 days (proteomics) in hypoxia (0.2% $O_2$) or normoxia (21% $O_2$). The heatmap shows common RNA and protein expression patterns between hypoxic and normoxic samples. Highest similarities in RNA and protein expression patterns were found in samples from the same cell line, highlighting cell line variability. Unsupervised clustering was performed by calculating Canberra distances of Z-scores for biological repeats. Each column within the heatmap represents a biological repeat, having three biological repeats per cell line and condition.

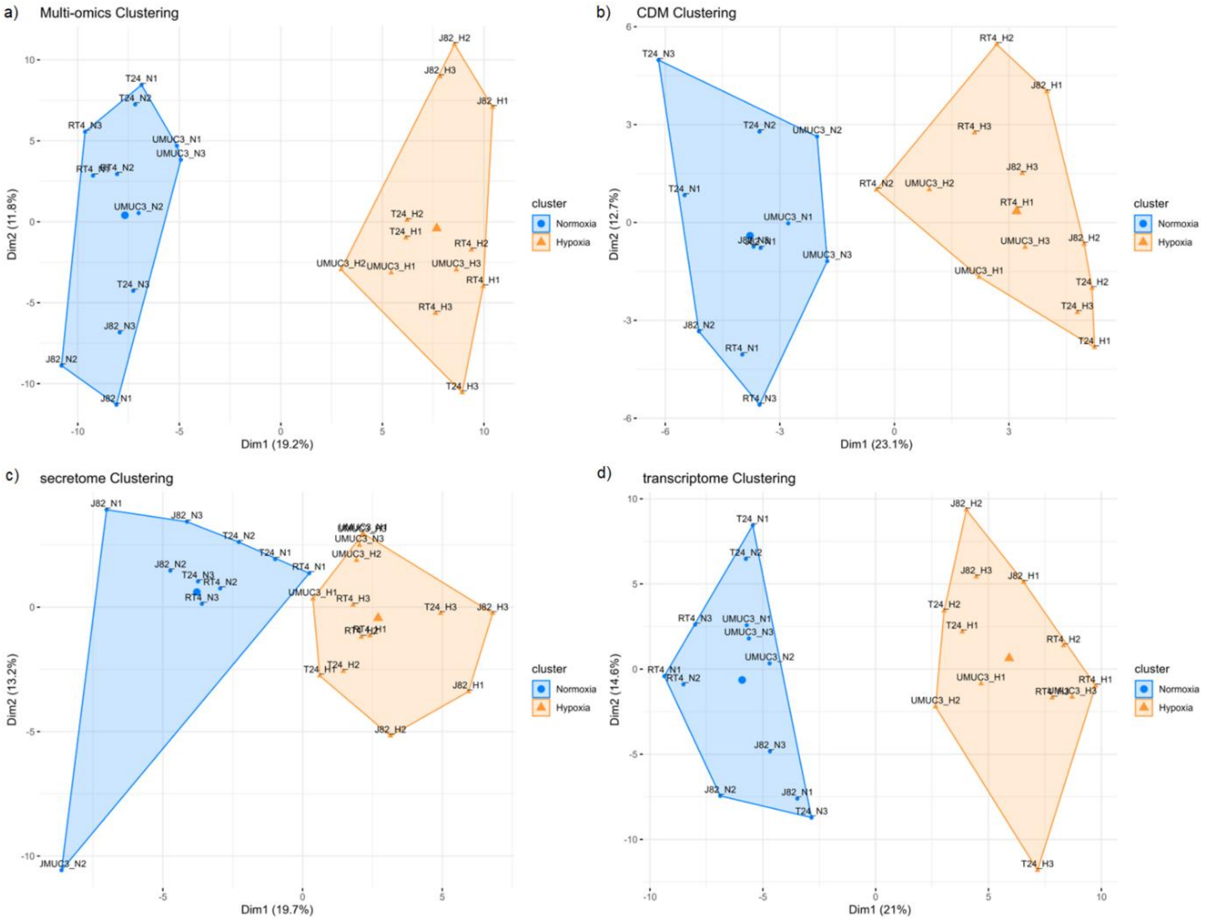


**Figure S3: K – means clustering suggests hypoxia induces common changes in ECM gene expression at protein and RNA levels.** Figure represents unsupervised k-means clustering for Z-scores from significant ($p_{adj}<0.05$, fold change ≥2 or ≤-2) RNA (n=196) and proteins (n=66 CDM, n=67 secretome). Clear separation between hypoxic (H) and normoxic (N) samples can be observed for the multi-omics (a) CDM (b), secretome (c), transcriptome (d) datasets. Unsupervised clustering was performed using either Euclidean (multi-omics, secretome) or Manhattan (CDM, transcriptome) distances of Z-scores for biological repeats. The optimal method to calculate distance was determined for each independent dataset using both Adjusted Rand Index and Silhouette methods. Analyses included three biological repeats per cell line and condition.

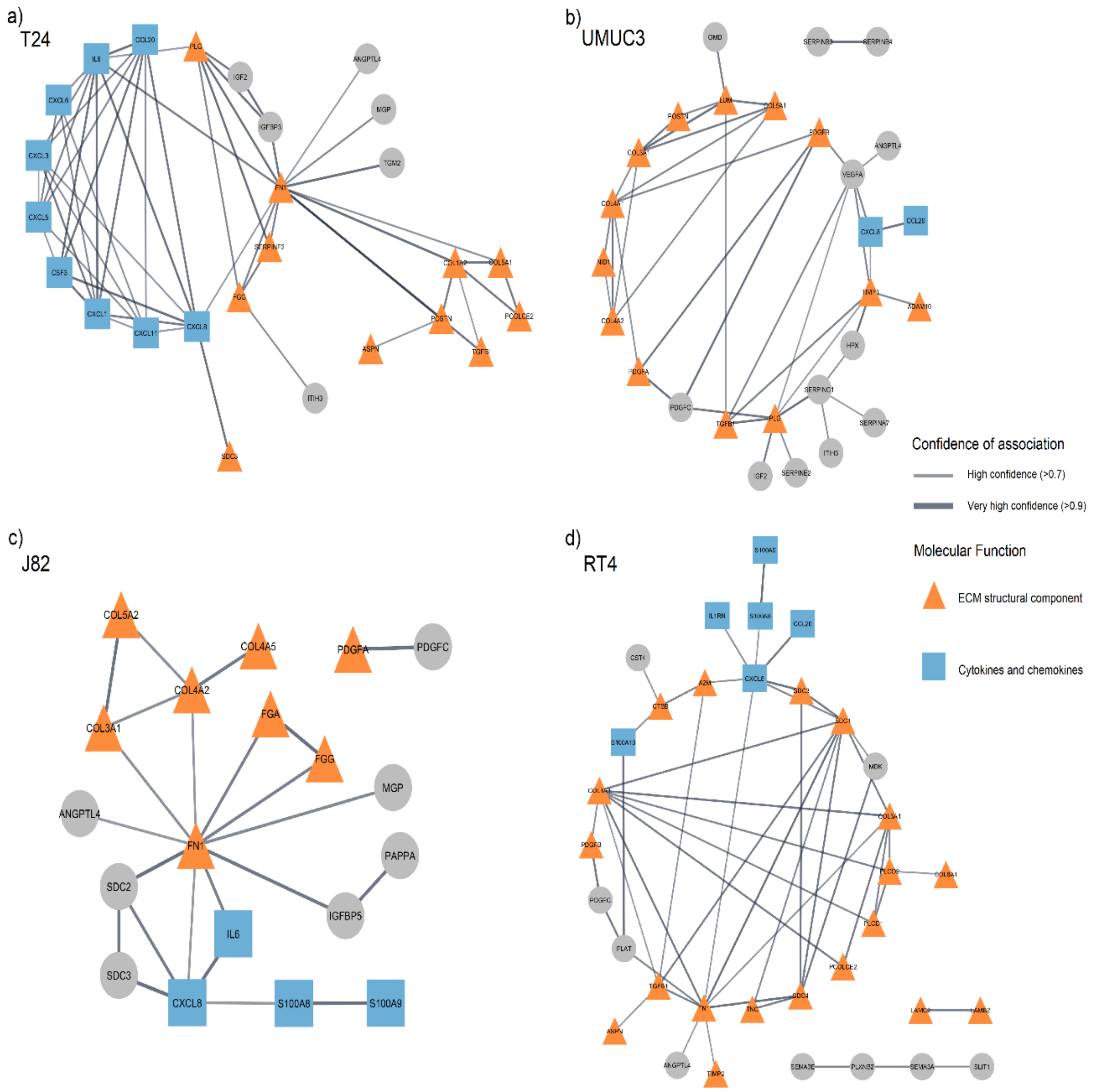


**Figure S4: Protein-protein interaction (PPI) networks predict strong interactions between structural and immune ECM proteins.** The figure represents the predicted protein interactions for those ECM protein with significant change ($p_{adj}\leq0.05$, fold change≥2) due to hypoxia stress for T24 (a), UMUC3 (b), RT4 (c) and J82 (d) cell lines in both CDM and secretome protein fractions. The PPI networks highlight a predominant role of fibronectin (FN) and collagen (COL) proteins, being key interacting nexus in all networks. Furthermore, strong interactions are predicted between structural ECM proteins and immune signalling proteins in all cell lines, but especially in T24 cells. The data suggests hypoxia induces ECM protein changes mostly affecting structural ECM protein interactions but also cytokines and chemokines interactions, predicting a strong link between both protein groups. Data were generated in three independent experiments per cell line. Confidence score represents the strength of the association (0-1 scale) according to the STRING Database costumed confidence score calculations.

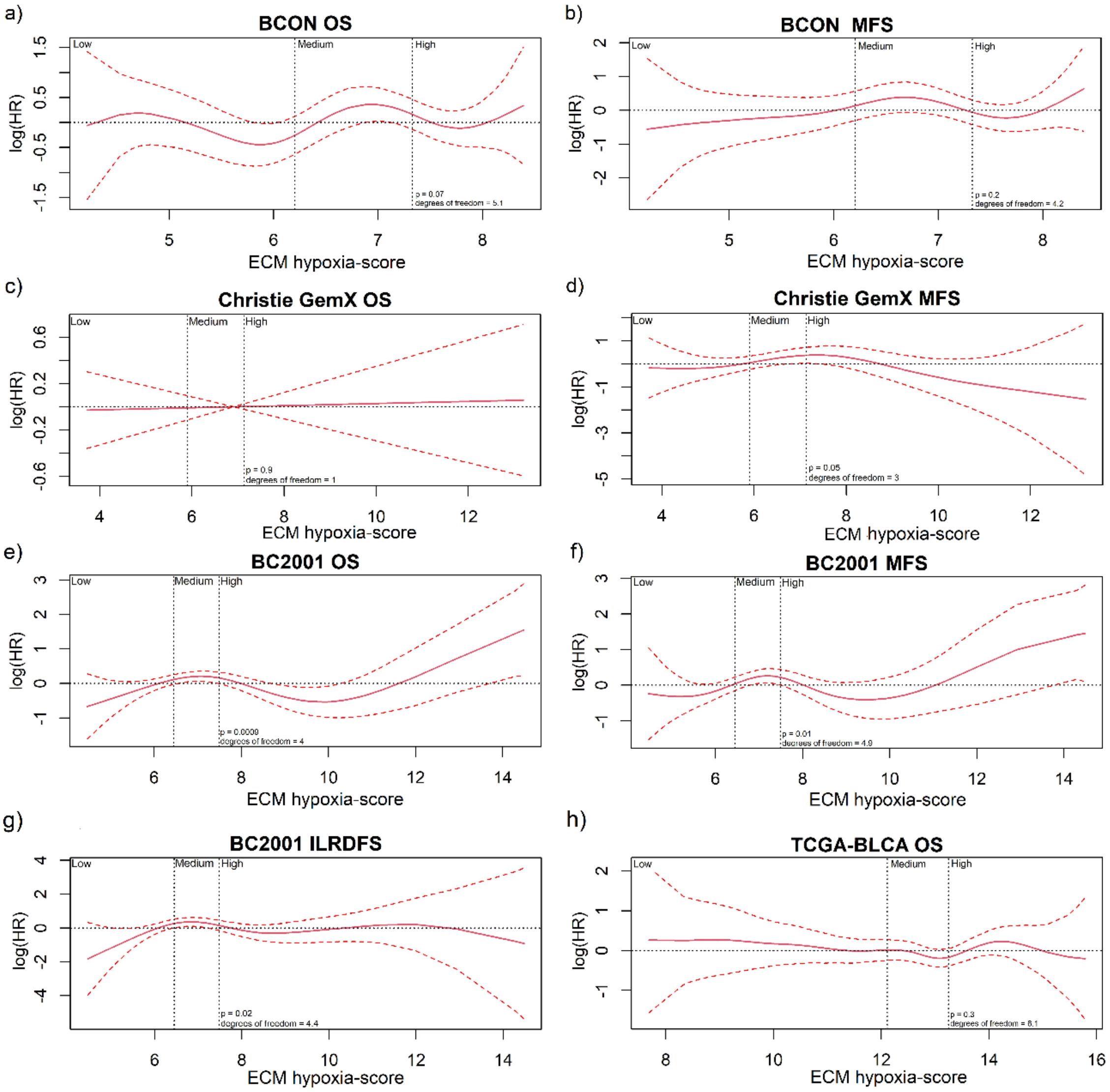


**Figure S5: A candidate hypoxia-derived ECM signature shows non-linear association patterns with hazard ratio (HRs).** A candidate ECM signature (*COL5A2, CSTB, CTSH, S100A2 and S100A9*) was retrospectively validated in the BCON (n=151; a, b), Christie GemX (n=183; c, d), BC2001 (n=313; e-g) and TCGA-BLCA (n=397; h) cohorts up 10 years of follow-up. Univariable Cox regression was used to evaluate the continuous relationship between ECM hypoxia-score with hazard ratios. Tertile stratification was used to identify the low, medium and high groups patient classification based on the ECM gene signature median expression levels. Cox regression models show a non-linear association pattern between ECM hypoxia-scores and HRs for all variables and cohorts with only two exceptions: Christie GemX overall survival (OS) (c), and TCGA-BLCA OS (h). Patients with medium ECM hypoxia-score values had higher HRs in the BCON (a, b), Christie GemX (d) and BC2001 (c-e) cohorts. Significant association between ECM hypoxia-score and HR was found in OS, metastasis-free survival (MFS) and invasive locoregional disease-free survival (ILRDFS) at the BC2001 cohort (c-e), and for MFS at the Christie GemX cohort (d). In BCON, Christie GemX and BC2001 cohorts, all patients underwent UK standard-of-care radiotherapy. In the TCGA-BLCA cohort, patients who underwent radiotherapy (n=12) were excluded, considering it a surgical cohort. n=1 and n=2 patients had not available overall survival data in the Christie GemX and the TCGA-BLCA cohorts, respectively. *S100A2* expression levels were not available in the BCON cohort and therefore it was not included in the signature. Analyses were performed retrospectively using Cox proportional hazards model.

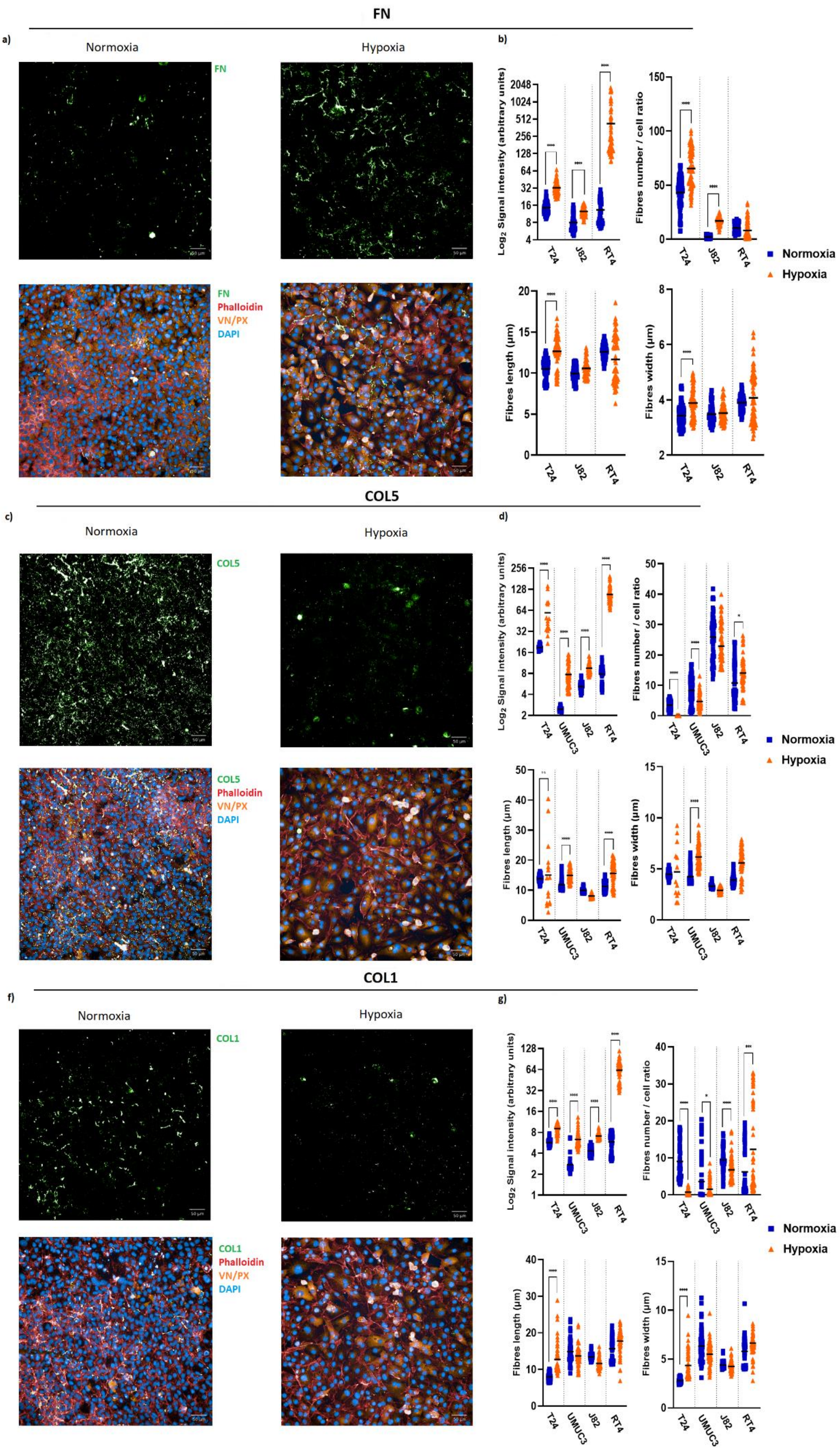


**Figure S6: Hypoxia affects the morphology and number of ECM fibres.** The figure shows immunofluorescence images of ECM produced under hypoxic stress by T24 cells stained for FN (a), COL5 (c) or COL1 (e) (green). Staining was also performed for VN/PX (yellow), actin (red) and nuclei (blue). Bar plots show fibre intensity, number, width and length for FN (b), COL5 (d) and COL1 (f). Analyses show a significant increase in signal intensity for FN, COL1 and COL5 in hypoxia. A significant increase in fibre numbers was seen for FN. However, COL1 and COL5 show a significant decrease in fibre number. Finally, a significant increase in fibre length is also seen for FN, while COL1 fibres had a significant increase in both fibres length and width. Significance is defined as p≤0.05 and at least 20% fold-change, with * for p≤0.05, ** for p≤0.01, *** for p≤0.001 and **** for p≤0.0001. Signal intensity was normalised to total cell number. Data are from 3 biological repeats, with 20 technical repeats for each biological repeat.

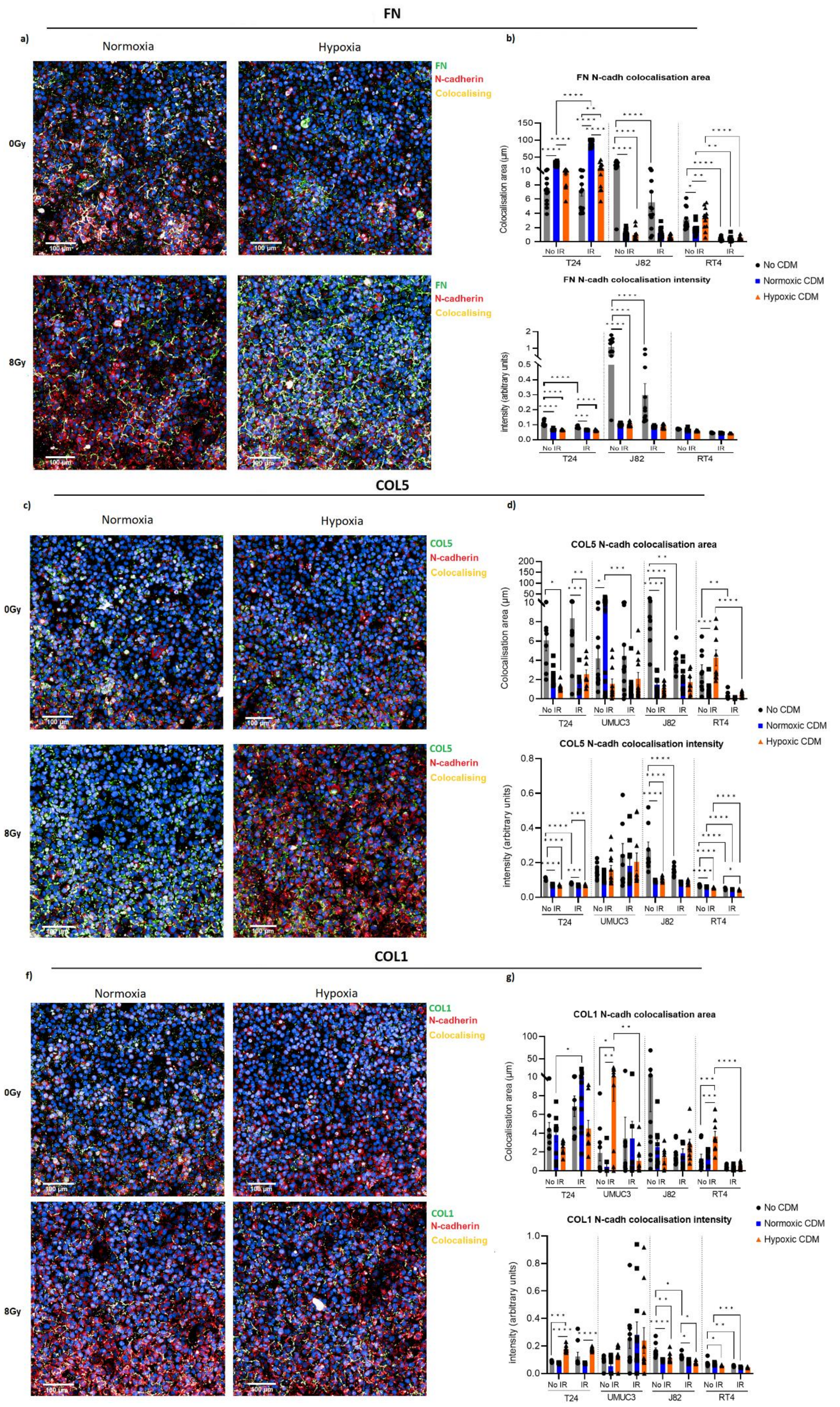


**Figure S7: Hypoxic ECM generally impairs N-cadherin co-localisation with FN and COL1 fibres.** The figure shows immunofluorescence images for FN (a), COL5 (c) or COL1 (e) produced in hypoxia (0.2% $O_2$) and normoxia (21% $O_2$) and their co-localisation with N-cadherin in non-irradiated (0Gy) and irradiated (8Gy) T24 cells. Bar plots show the area and intensity of N-cadherin co-localised with FN (b), COL5 (d) and COL1 (g) fibres in T24, UMUC3, J82 and RT4 cells. Non-coated controls were included. No FN synthesis was detected for UMUC3, and it was not analysed for FN co-localisation. Analyses show a significant decrease in N-cadherin co-localisation in T24, but increase in RT4, for hypoxic vs normoxic FN fibres for non-irradiated (T24, RT4) and irradiated (T24) cells. Decreased N-cadherin co-localisation area was also seen in UMUC3 and RT4 with hypoxic vs normoxic COL1 fibres for non-irradiated cells. Increased intensity of co-localised N-cadherin was seen in T24 with hypoxic vs normoxic COL1 fibres, independently of irradiation. After irradiation, a significant decrease in co-localised E-cadherin area was seen with FN J82, RT4), COL5 (UMUC3, RT4) and COL1 (UMUC3, RT4). However, N-cadherin co-localisation area increased with FN and COL1 after irradiation in T24. Co-localised N-cadherin intensity was also reduced after irradiation in RT4 for FN and COL5 and COL1 fibres. Significance was measured with 2-way ANOVA with Bonferroni correction, being as $p<0.05$ and at least 20% fold-change, with * for $p<0.05$, ** for $p<0.01$, *** for $p<0.001$ and **** for $p<0.0001$. Signal intensity and colocalisation area was normalised to total cell number. Data are for 3 biological repeats with at least 3 technical repeats for each biological repeat.

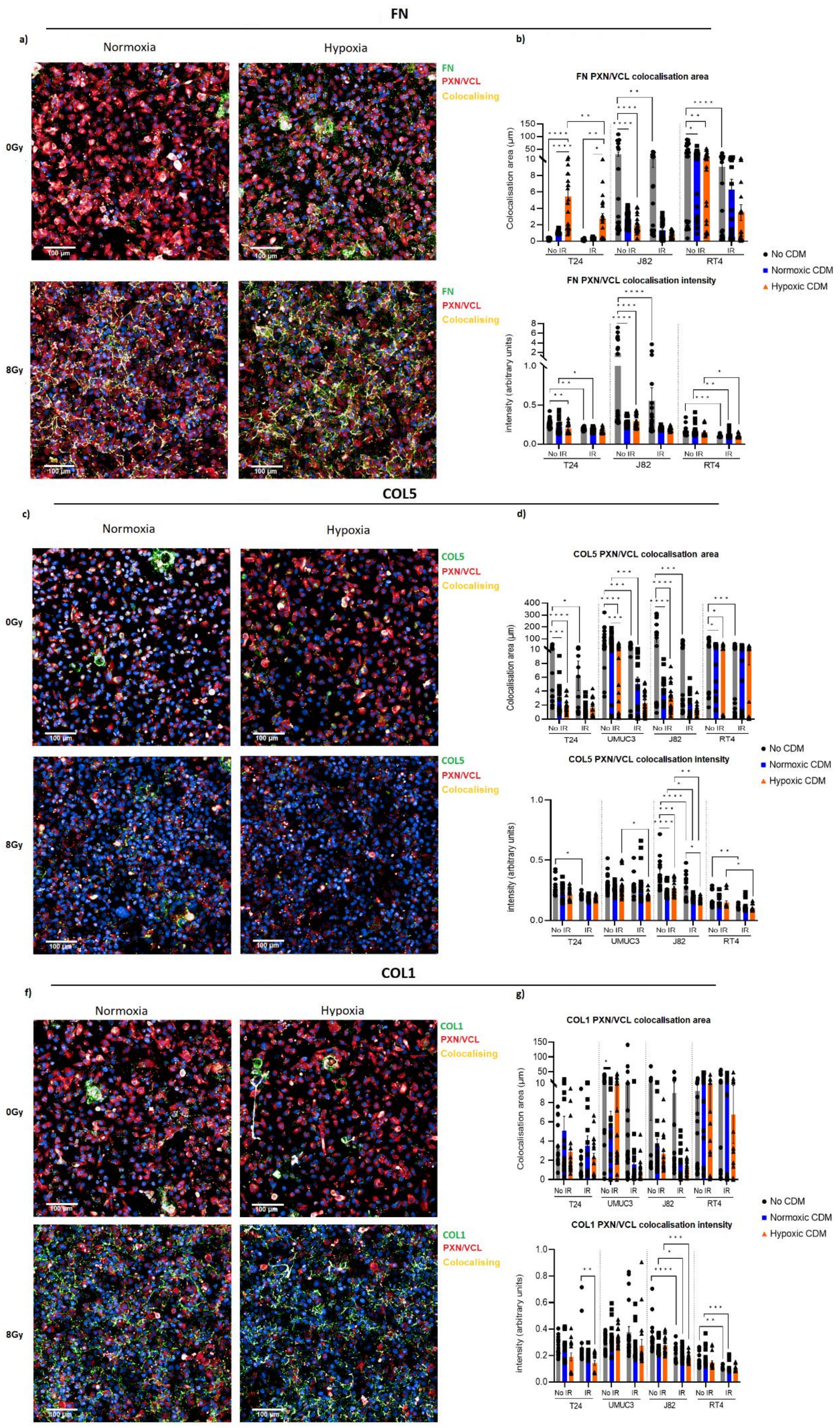


**Figure S8: Irradiation (IR) generally impairs paxillin/vinculin (PX/VN) co-localisation with ECM fibres.** The figure shows immunofluorescence images for FN (a), COL5 (c) or COL1 (e) produced in hypoxia (0.2% $O_2$) and normoxia (21% $O_2$) and their co-localisation with PX/VN in non-IR (0Gy) and IR (8Gy) T24 cells. Bar plots represent area and intensity of PX/VN co-localised with FN (b), COL5 (d) and COL1 (g) fibres in T24, UMUC3, J82 and RT4 cell line. Non-coated control was included. No FN synthesis was detected for UMUC3, and therefore was excluded from FN co-localisation analysis. Analysis showed significant increased PX/VN co-localisation area in T24 with FN, but decreased PX/VN co-localisation area for UMUC3 with COL5. Hypoxic ECM fibres did not affect PX/VN co-localisation intensity. After IR, PX/VN co-localisation area was reduced for T24 with FN, and for UMUC3 with COL5. Colocalised PX/VN intensity was also reduced for T24 and RT4 for FN, for UMUC3, J82 and RT4 for COL5, and J82 and RT4 for COL1. Significance was measured with 2-way ANOVA with Bonferroni correction, being defined as $p<0.05$ and at least 20% fold-change, with * for $p<0.05$, ** for $p<0.01$, *** for $p<0.001$ and **** for $p<0.0001$. Signal intensity and colocalisation area was normalised to total cell number. Data are for 3 biological repeats with at least 3 technical repeats for each biological repeat.

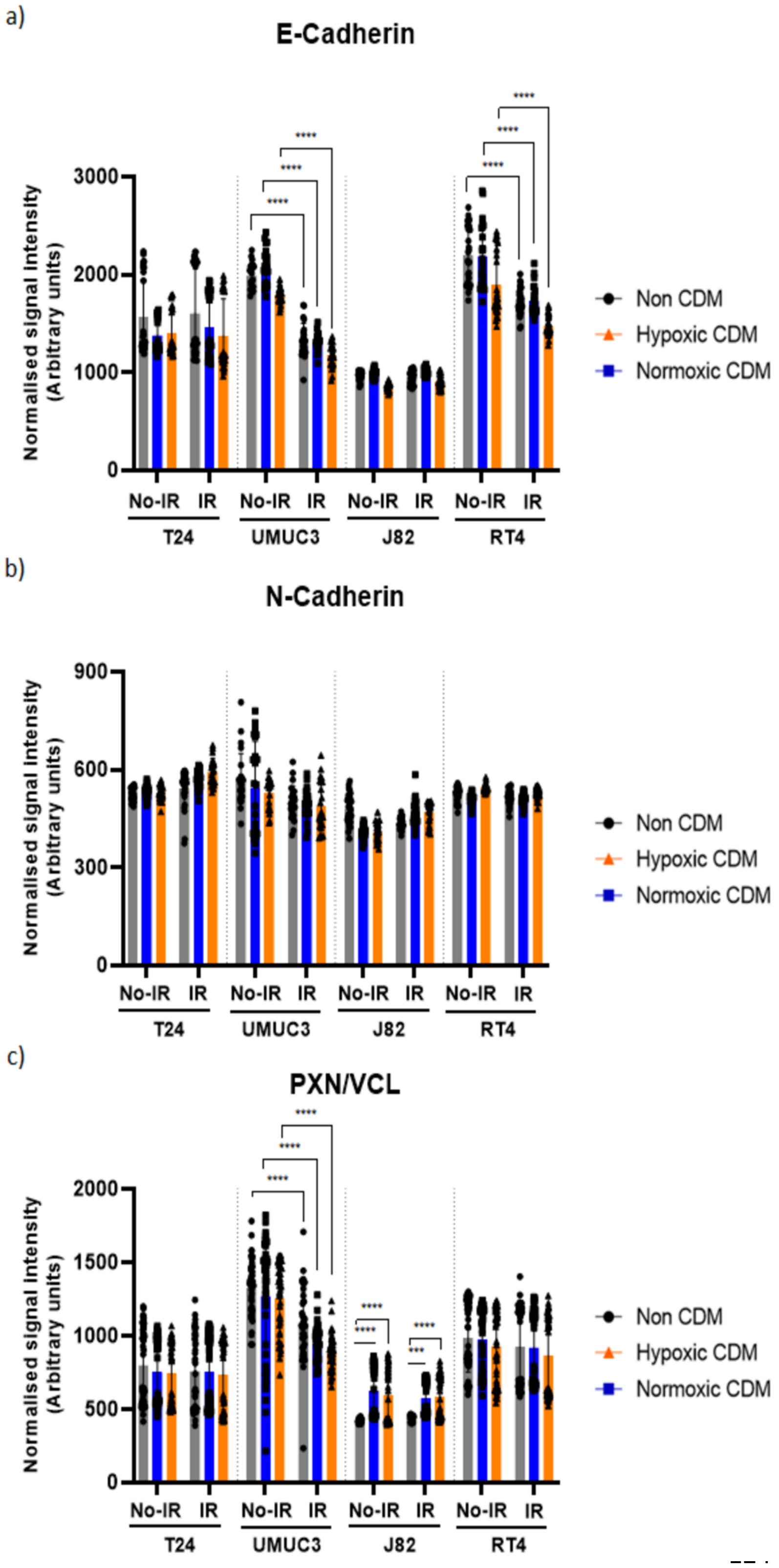


**Figure S9: Irradiation (IR) reduces total expression levels of E-cadherin and paxillin/vinculin (PXN/VCL).** Barplots represent the normalised signal intensity of E-cadherin (a), N-cadherin (b) and paxillin/vinculin (PXN/VCL) (c) in T24, UMUC3, J82 and RT4 cell line. Non-coated control was included. Analysis showed significant decreased normalised signal intensity for E-cadherin (UMUC3, RT4), and paxillin/vinculin (UMUC3), independently of cells being seeded on top of hypoxic or normoxic CDMs. No significant changes were observed for N-cadherin. Cell seeding onto hypoxic or normoxic CDM coating did not affected E/N-cadherin or PXN/VCL levels. Results suggest IR can affect E-cadherin and PXN. Significance is defined as p<0.05 and at least 20% fold-change, with * for p<0.05, ** for p<0.01, *** for p<0.001 and **** for p<0.0001. For each experiment, n=3 biological repeats, with at least n=3 technical repeats for each biological repeat.

### Supplementary methods

**Christie GemX cohort:** Christie GemX (09/H1013/24) is a muscle-invasive bladder cancer (T2–T4, N0, M0) cohort (n=184). Patients were diagnosed between 2010 and 2017, median follow-up was 78 months (range: 17-165 months), and treatment followed the current UK standard-of-care radiotherapy (55 Gy in 20 fractions over 4 weeks). Patients received gemcitabine (100 mg/m$^2$ once a week) (median 3 cycles; range 1-4), and 111 patients received neoadjuvant chemotherapy (96 cisplatin and gemcitabine + GemX, 17 carboplatin and gemcitabine + GemX, 6 carboplatin and etoposide + GemX). One patient had no outcome data available. Transcriptomic data was generated using a previously described analysis pipeline[20].

**Cell culture.** Bladder cancer cell lines (T24, UMUC3, J82, RT4) were acquired from the American Type Culture Collection (ATCC; Virginia, USA). Cells were routinely authenticated and tested for mycoplasma. Cells were cultured in McCoy's modified 5A medium with L-glutamine (Gibco, Waltham, USA) supplemented with 10% foetal bovine serum (FBS) (Sigma-Aldrich, Missouri, USA). A hypoxia station (Don Whitley Scientific Bingley, UK) (37ºC, 5% $CO_2$) was used to culture cells at 1%, 0.2%, or 0.1% $O_2$ either continuously (chronic) or in 24 h cycles of hypoxia (transient) for 1, 2, 4, or 7 days. Control cells were cultured in parallel under normoxic conditions (20.9% $O_2$, 37°C, 5% $CO_2$).

**Meta-analysis of transcriptomic data:** Genes with tau-squared >0.1 were discarded to avoid data biases due to datasets differences. Percentage frequency, defined as the number of times a gene is identified as significant across all meta-analyses datasets groups and expressed in 0-1 scale, was calculated for each meta-analysis gene. An expression score was calculated to account for the frequency a gene was identified as significant, defined by the following formula:

$$expression_{score} = (-log10(FDRrem) \times logFCrem) \times percentage_{frequency}$$

For all further analyses, significance was defined as percentage_frequency >0.2, $FDR_{rem}<0.05$, and $FC_{rem}>1$ or $FC_{rem}<-1$. Z-scores were calculated using gene scores as reference. Similarity scores were calculated independently for each gene using cosine similarity distances scaled from 0 to 1 as per the following formula:

$$scaled = \frac{\left(\frac{\sum(gene_{score} \times ref_{value})}{\sqrt{\sum(gene_{values}2)} \times \sqrt{\sum(ref_{value}2)}}\right) + 1}{2}$$

In which ref_values represents the calculated gene score at pan-cancer level, which was used as reference, whilst expression_score represents the previously calculated expression score for cancer REM dataset. Average scaled similarity score across all genes was then used to determine the similarity level of each cancer type with the pan-cancer dataset. Ontology enrichment analyses were performed using the *clusterProfiler*, *ReactomePA*, *AnnotationDbi* and *org.Hs.eg.db* packages. Ontology, general data processing and manipulation, and figures representation was performed in R (v.4.02 – v.4.4.2) using the packages *dplyr*, *tidyr*, *matrixStats*, *ComplexHeatmap*, *circlize*, *gridExtra*, *RColorBrewer*.

**Production and quantification of cell lysates, cell-derived extracellular matrices (CDMs) and secretomes:** Whole-cell lysates were obtained by scraping with Pierce IP Lysis Buffer (Thermo Fisher Scientific, Massachusetts, USA). Samples were sonicated using a Bioruptor Pico sonicator (Diagenode New Jersey, USA) and centrifuged (10 min, 21000 rcf, 4°C) to remove debris.

CDMs were extracted as previously described[70]. Briefly, cell decellularisation was performed using extraction buffer (20 mM $H_4OH$, 0.5% Triton X-100 in phosphate buffered saline [PBS]). Residual DNA was removed by incubation with DNase I (Invitrogen, Massachusetts, USA) solution (20 U/ml in PBS, 30 min, 37°C). CDM was recovered by scraping with 2X SDS buffer (4% (w/v) SDS, 10% (w/v) glycerol, 50 mM Tris HCl, 0.005% (w/v) bromophenol blue, 20% (v/v) mercaptoethanol).

Secretomes were extracted as previously described[70]. Briefly, cells were grown for 4 days in hypoxia with serum-supplemented medium, and 3 days in serum-free medium. Serum-free media were collected and concentrated in a 10-10000 Da MWCO centrifugal concentrator (Sartorius, Göttingen, Germany; 2 h, 4000 rcf, 4°C).

Whole-cell lysate and secretome protein concentrations were measured using a Pierce BCA Protein Assay Kit (Thermo Fisher Scientific) following manufacturer's instructions. CDM protein concentration was determined using InstantBlue (Abcprovideam, Cambridge, UK) after SDS-PAGE electrophoresis (45 min, 200 V; 4-12% Bis-Tris gels; Thermo Fisher Scientific), using protein samples of known concentrations as standards as previously described[71].

**Western blotting.** Whole-cell lysates (50 µg) and CDM (10 µg) samples were used for western blotting for GLUT1 and FN, respectively. Samples were separated by SDS-PAGE electrophoresis (45 min, 200V; 4-12% Bis-Tris gels; Thermo Fisher Scientific) using NuPAGE MES SDS Running Buffer (Thermo Fisher Scientific). Transfer (10 h, 30 V, 4°C) to nitrocellulose membranes (Whatman, Maidstone, UK) was performed with 1X transfer buffer (25 mM Tris, 192 mM glycine 20% methanol [v/v]). Membranes were blocked (1 h, room temperature) with casein-blocking buffer (Sigma-Aldrich; 1:10 dilution) in TBST (10 mM Tris-HCl, pH 7.4, 150 mM NaCl, 0.05% [w/v] Tween-20). Membranes were stained with Ponceau solution (Thermo Fisher Scientific) and imaged according to the manufacturer's protocol. Membranes were then incubated with anti-GLUT1 (Merck Millipore,

Massachusetts, USA; 1:1000 dilution) or anti-FN (Sigma-Aldrich; 1:1000 dilution) rabbit polyclonal antibodies. Anti-cyclophilin-β rabbit polyclonal antibody (Abcam; 1:6000 dilution) was used as a loading control, and incubations were performed in TBST-casein blocking buffer overnight at 4°C. Membranes were then incubated with polyclonal goat anti-rabbit secondary Alexa Fluor 680 nm (Thermo Fisher Scientific; 1:10000 dilution; 2 h, room temperature, dark conditions). Imaging was performed using an Odyssey Infrared Imaging System (LI-COR Biosciences, Lincoln, USA) at 700 nm. Band signal intensity was determined based on the median pixel signal intensity using Odyssey software (LI-COR Biosciences).

**Mass spectrometry (MS).** CDM samples were separated using SDS-PAGE (3 min, 200V; 4-12% Bis-Tris gel; Thermo Fisher Scientific). Protein bands were stained with InstantBlue Coomassie (Abcam), processed for in-gel trypsin digestion, and analysed by tandem LC-MS/MS using an UltiMate 3000 Rapid Separation LC system (Thermo Fisher Scientific) coupled with an Orbitrap Elite Mass Spectrometer (Thermo Fisher Scientific) following previously described protocols[9,70].

**MS data processing.** MS data were analysed using an in-house Mascot Server (v. 2.5.1; Matrix Sciences) as described previously[9,72], with mass tolerances of 0.4 Da (precursor ions) and 0.5 Da (fragment ions). Data were validated in Scaffold (v. 4.6.3; Proteome Software) using a threshold of identification of 90% at the peptide level, 99% at the protein level, and with at least one unique validated peptide (0.1% of estimated false discovery rate with these settings). MS protein identification, total protein normalisation, fold-change and p-values were calculated using Protein Discoverer (v. 2.3.0.523).

**Transcriptomics.** Cells were cultured for 24 h in 0.2% $O_2$, extracting RNA (RNeasy Plus Mini Kit; Qiagen, Hilden, Germany), and hybridising onto Clariom S Pico HT human gene expression arrays (Thermo Fisher Scientific). Gene expression data were normalised using the apt-probeset-summarize application (v1.20.0; Thermo Fisher Scientific), applying GC content and RMA-style background corrections (option: gc-sst-rma-sketch). Resulting expression matrices were corrected for batch effects using either sva:ComBat (v3.50.0)[73] or limma:removeBatchEffect (v3.60.3)[74]. Fold changes (FCs), p-values, adjusted p-values and $log_2$ normalisation (rlog function) were calculated using *limma* (v3.60.3)[32] and *DESeq2*(v.1.42.0)[31]. All analysis were performed in R (v.4.2.2).

**ChIP-seq.** ChIP-seq was performed as described previously[53]. Briefly, T24 cells were cultured for 24 h in 0.1% $O_2$. The protein-DNA interactions were cross-linked using ChIP cross-linked gold (Diagenode, New Jersey, USA; 10 min, room temperature) and 1% formaldehyde (10 min, room temperature) before lysing cells and shearing chromatin into 200-300bp fragments using a Bioruptor Pico

(Diagenode). Immunoprecipitation was induced by incubating overnight at 4°C with antibodies against HIF-1α (Abcam; 1 mg/ml), HIF-1β (Novus Biological, Colorado, USA; 1 mg/ml), HIF-2α (Abcam; 1 mg/ml), and Dynabeads Protein G (Thermo Fisher Scientific) in blocking buffer (5 mg/ml bovine serum albumin [BSA] in PBS). Precipitated fragments were de-cross-linked, and DNA was eluted using phenol-chloroform. Sequencing and mapping were performed using the CRUK-MI core facilities.

**Multi-omics *in silico* analysis.** Omics data was integrated using the *metafor:rma* function[34]. Gene similarity levels were determined as per the hypoxia transcriptomic meta-analysis. Clustering analyses were performed using the original omics *in vitro* datasets for genes significant in at least one cell line ($p_{adj}$<0.05, FC>2 or FC<-2). Multi-omics clustering was performed by combining all omics datasets. ChIPSeq dataset was used to annotate HIF1/2 genes regulated genes. ECM gene ontologies were annotated using The Matrisome Project Database[35]. Ontology, clustering analyses, and figure representation were performed using R (v.4.02 – v.4.4.2) packages: *org.Hs.eg.db*, *AnnotationDbi*, *ClusterProfiler*, *DOSE*, *ggplot2*, *enrichplot; ReactomePA; GoSemSIM; factoextra; cluster; mclust; ggVennDiagram*. Protein-protein interaction (PPI) analysis was performed for ECM genes significant at protein level using Cystoscope (v.3.9.1; National Institutes of Health, Washington, USA)[75], with confidence scores (0-1 scale) calculated using the STRING database (v.11.5; University of Zurich, Zurich, Switzerland)[76].

**Definition of outcomes for clinical cohorts:** The co-primary endpoints were overall survival (OS), defined as the time from the date of randomisation to the date of death due to any cause. Secondary endpoints were cancer specific survival (CSS), defined as the time from the date of randomisation to the date of death due to bladder cancer, disease-free survival (DFS), defined as from the date of randomisation to cancer recurrence or death from any cause, metastasis-free survival (MFS), defined as the time from the date of randomisation to the date of diagnosis of metastasis, recurrence free survival (RFS), defined as from the date of randomisation to the date of disease recurrence, progression-free survival, defined as from the date of randomisation to cancer recurrence, metastasis, or death from any cause, invasive locoregional disease-free survival (ILRDFS), defined as from the date of randomisation to the date of invasive disease recurrence, locoregional disease-free survival (LRDFS), defined as from the date of randomisation to the date of non-invasive disease recurrence, and for prostate cancer, biochemical recurrence (BCR), defined as from the date of randomisation to the date of biochemical disease recurrence (prostate specific antigen level increase). Predictive capacity (radiotherapy versus surgery) of the signature was performed by meta-analysing datasets (bladder and head & neck cancer). For breast, cervix and pancreatic, only TCGA data was used. For glioblastoma, only CGGA data was used.

**Artificial Neural Network (ANN) modelling:** A combined MIBC cohort was divided into training (80%) and validation (20%) sets. Treatment (radiotherapy, cystectomy) and ECM score groups (low & high, medium) were used as input variables, with OS as output variable. Data augmentation was performed via random shuffling of input values. Artificial values (n=197) were added to the original dataset of "alive" patients to correct the imbalance between those who died (n=728) and those who survived (n=531). Data augmentation has been used previously in clinical prediction models to avoid biased predictions and improve robustness[77].

ANN followed an analysis approach with a feedforward structure tailored for binary classification tasks. Two hidden layers and one output layer were used. The first layer used 24 neurons, whereas the second layer used six neurons, both activated by the rectified linear unit (ReLU). To regulate overfitting, a neuron deactivation step follows, randomly deactivating 10% of the neurons during training in both layers independently. In addition, L2 Kernel regularisation was applied to the first hidden layer by adding a penalty charge to the loss function during training. The output layer employed a sigmoid activation function to calculate probabilities indicating binary class predictions. Optimisation was facilitated using the Adam optimiser, with accuracy serving as the main metric for performance assessment.

A Receiver Operating Characteristic curve was used to estimate true vs. false positive prediction rates, calculated as Area Under the Curve (AUC) values. Prediction model and all associated analyses were performed using Python v.3.12.2 (Python Software Foundation, DE, USA) with the following packages: *pandas*, *numpy*, *tensorflow.keras*, *matplotlib.pyplot*, *seaborn*, *scikit-learn*, *keras.regularizers*, *sklearn.preprocessing*.

**Cell seeding onto CDM coatings.** Cells were grown in normoxia (20.9% $O_2$) or hypoxia (0.2% $O_2$). CDM coatings were produced by plate decellularisation with $NH_4OH$ lysis buffer. Control plates without CDM were "mock" decellularised. CDM was washed with PBS, blocked for 1 h at room temperature (10% w/v bovine-serum albumin in PBS), and immediately used for cell culture. Cells were irradiated (0-8 Gy) using an Xstrahl CIX3 irradiator (Xstrahl, Camberley, UK), and seeded onto CDM coatings produced by the same cell line.

**Attachment and migration assays:** To assess migration, cells were seeded onto CDMs to achieve 100% confluence: $2x10^5$ cell/ml (0 Gy); $3x10^5$ cell/ml (2 Gy); $4x10^5$ cell/ml (4 Gy); $6x10^5$ cell/ml (6 Gy), or $1x10^6$ cell/ml (8 Gy). After 24 h at 37°C cultures were scratched using a WoundMaker System (Essen Biosciences, Michigan, USA). Cell migration was measured by acquiring images (one picture/h) at 20X magnification per well over a period of 24 h (T24) or 48 h (UMUC3, J82) using the

IncuCyte S3 Microscopy System (Essen Biosciences). Cells were maintained at 37°C throughout. Images were analysed using the IncuCyte Scratch Wound Analysis Software (Essen Biosciences).

To assess attachment, new cells (16,000 cells/well) were seeded and allowed to attach for 30 min (T24, RT4, J82) or 1 h (UMUC3) at 37°C. Unattached cells were removed, and the remaining cells were fixed with 5% glutaraldehyde. Fixed cells were stained with 0.1% (v/v) crystal violet solution in 200 mM MES buffer (pH=6.0) for 60 min at room temperature. Crystal violet was solubilized with 10% acetic acid (v/v), and absorbance was measured at 570 nm with a Varioscan LUX plate reader (Thermo Fisher Scientific).

**Immunofluorescence, co-localisation and image analysis:** Three-step sequential staining was performed as follows: (1) blocking with 1% (w/v in PBS) BSA (1 h, room temperature), followed by overnight incubation (4°C) with rabbit polyclonal antibodies for FN (Sigma-Aldrich, 1:300 dilution), COL5 (Novus Biological), or COL1 (Novus Biological) in 1% (w/v in PBS) BSA blocking buffer. (2) permeabilisation with 0.3% (v/v) Triton-100X in PBS (30 min, room temperature), followed by overnight incubation (4°C) with VCL (Sigma-Aldrich, 1/300 dil.) and PXN (BD Biosciences, New Jersey, USA; 1/300 dilution) monoclonal mouse antibodies (Thermo Fisher Scientific) in 1% (w/v in PBS) BSA blocking buffer. (3) incubation with 488 nm Alexa Fluor anti-rabbit (Thermo Fisher Scientific; 1/500 dilution), Alexa Fluor 546 nm anti-mouse (1/500 dilution) secondary antibodies, and Alexa Fluor 647 nm phalloidin (1/400) in 1% (w/v in PBS) BSA blocking buffer (2 h, room temperature). Nuclei were stained with 300 mM DAPI (5 min, room temperature). Samples were imaged using high content screening with PerkinElmer Operetta (PerkinElmer), a wide-field fully automated high-content screening system with a range of eight LEDs for excitation of fluorophores (365, 405, 440, 475, 510, 550, 630, and 660 nm) and a range of emission filters (430-500, 460-515, 470-515, 500-550, 570-580, 655–705 nm). Digitisation was performed with a Zyla sCMOS camera (2160 × 2160 pixels, 6.5um pixel size, 16-bit, readout noise <1.2e; Andor, Belfast, UK).

For co-localisation experiments, cells were seeded onto CDMs coatings. After FN1, COL5, and COL1 antibody incubation, samples were incubated with 488 nm Alexa Fluor anti-rabbit in 1% [w/v] BSA (2h, room temperature). Subsequent overnight incubation was performed with E-cadherin polyclonal rabbit antibody (Proteintech, Manchester, UK) or N-cadherin mouse polyclonal antibody (Abcam), in addition to VN and PX antibodies. Samples were incubated (1% [w/v] BSA, 2h at room temperature) with anti-mouse Alexa Fluor 647 nm (Thermo Fisher Scientific) and Alexa Fluor anti-rabbit 546 nm (Thermo Fisher Scientific). Samples were imaged using high-content screening via PerkinElmer Opera Phenix (PerkinElmer), a confocal spinning disk 4 laser (405 nm 50mW, 488 nm 50mW, 561 nm 50mW, 640 nm 50mW) fixed light path system, with a range of emission filters (435–

550 nm, 435–480 nm, 500–550 nm, 570–630 nm, 650–760 nm). Four Zyla sCMOS cameras, 2160x2160 pixels, 6.5um pixel size (Andor, Belfast, UK) were set up for each dedicated light path. 20 fields of view were acquired with a Z range of 34.5µm in 1.5µm steps using the Zeiss W Plan-Apochromat X20 water objective NA 1.0 WD 1.17 mm.

**Statistical analysis of *in vitro* mechanistic data.** Immunofluorescence and attachment data normality were tested with Shapiro-Wilks. Statistical differences were calculated using unpaired t-tests or Mann-Whitney U-tests. Scratch assay statistical differences were determined with one-way variance analysis with Geisser-Greenhouse correction. Immunofluorescence co-localisation data was analysed using two-way ANOVA with Tukey correction. Analyses were performed using GraphPad Prism (v. 9.3.1; GraphPad, Massachusetts, USA).
